# VigExp: A functionally verified platform for aiding cowpea (*Vigna unguiculata*) and related legume crop improvement

**DOI:** 10.64898/2026.06.30.735734

**Authors:** Huanan Su, Danielle Mazurkiewicz, Nial Gursanscky, Matteo Riboni, Martina Juranić, Susan D. Johnson, Jee Hao Yow, Jasmin Deo, Yuhan Liu, Alexandria Mattinson, Gloria León-Martínez, Rocio Escobar-Guzmán, Rigel Salinas-Gamboa, Itzel Amasende-Morales, Jean-Philippe Vielle-Calzada, Anna M. G. Koltunow, Brett J. Ferguson

## Abstract

Legumes include some of the world’s most significant crop species, such as cowpea (*Vigna unguiculata*), a subsistence crop widely grown in sub-Saharan Africa. Despite their importance, legume crop improvement is hindered by a lack of high-resolution expression data, particularly for reproductive tissues and cell types. Here, we report on VigExp, a tool for visualising cowpea gene expression datasets. We demonstrate its utility across a range of vegetative and reproductive cell types of varieties IT97K-499-35 and IT86D-1010, which exhibit 93.75% protein sequence conservation and are amenable to stable transformation. This includes previously published transcriptomes of vegetative, floral and seed tissues, combined with developmentally staged male and female reproductive tissues. Also integrated are novel transcriptomes of laser-captured cell types covering reproductive development from meiosis to early embryo formation post-fertilisation. Spatial expression patterns and transcript levels can be visualised through an electronic fluorescent pictograph (eFP) browser. Validated by RT-qPCR, *in situ* hybridisation, transgenic, and CRISPR gene editing analyses, the predictive accuracy of VigExp matches prior cowpea functional study observations. Critical genes for nodule development and regulation were also identified and their expression patterns established in cowpea. Novel reference genes, constitutively expressed gene promoters for visualisation markers/gene-editing, and tissue- and cell-specific gene promoters for targeting these regions, were identified. The A-type cyclin, *VuTAM2*, was also identified, with a critical role in male meiosis established. Collectively, VigExp represents an adaptable and updatable resource to support crop improvement in cowpea and other legumes, which are often highly syntenic with respect to genome composition.

## INTRODUCTION

Legumes are members of the phenotypically diverse Fabaceae family, which includes over 19,000 species, representing the third largest family of land plants. Many are agriculturally and economically significant as key food, feed, forage and pasture species. They offer an economical source of nutrients, containing high levels of protein, fibre, vitamins and minerals, providing considerable benefit to regions suffering from food insecurity (Foyer et al., 2016). They also promote soil health through their symbiotic relationship with nitrogen-fixing soil bacteria, called rhizobia, that convert atmospheric di-nitrogen into forms of nitrogen the host plant can use (Ferguson et al., 2010, 2019). As a result, legume plants can enhance agricultural sustainability, often used in intercropping and rotation programs to enrich soil conditions to improve the growth and yield of co-planted non-legume species (Crews and Peoples, 2004).

Cowpea (*Vigna unguiculata*) is a subsistence legume crop used for both forage and food. It was domesticated in sub-Saharan Africa, where it is estimated to feed over 200 million people (Carvalho et al., 2017), but it is not significantly traded outside of Africa. Despite the considerable importance of cowpea for food security, limited improvements have been made to existing varieties using traditional breeding approaches, which are often constrained by low genetic diversity (Boukar et al., 2016; Horn and Shimelis, 2020). This is due in part to the high degree of near-obligate self-pollination, which promotes inbreeding, a characteristic feature of many crop legumes. Cowpea is highly syntenic with other *Vigna* species and agriculturally important legume crops, such as common bean and soybean (Sprent, 2007; Supplemental Figure 1, phylogenomic tree generated by using IQ-TREE).

One approach to enhance breeding efforts is to identify genes and molecular pathways responsible for controlling traits of interest and subsequently target favourable alleles and genetic variation to develop superior new varieties. Biotechnology can be deployed to alter endogenous gene expression by loss- or gain-of-function via gene editing or insertion of favourable genes using genetic modification to introduce desirable traits. For example, recent introduction of Bt (*Bacillus thuringiensis*) Cry protein-encoding genes into cowpea has provided resistance to the devastating pest *Maruca vitrata* that can reduce crop yields by 80% (Bett et al., 2017, 2019; Mohammed et al., 2014; Addae et al., 2020; Nboyine et al., 2024). These transgenic cowpeas are currently grown and consumed in Nigeria (Umar et al., 2022; Abubakar et al., 2023).

Given the importance of cowpea and other legumes as food sources, attention has focused on genetic aspects of post-fertilisation seed formation, seed content and seed quality (e.g. Tayade et al., 2023). An improved understanding of reproductive development is critical for increasing legume seed yields and enhancing food security. Little is known about the molecular mechanisms driving early reproductive development leading to fertilisation and subsequent regulation of post-fertilisation seed development across the legumes in general (Erfatpour et al., 2024). There is also a lack of supporting transcriptomic and bioinformatic resources available that provide gene expression profiles during the developmental sequence of events in male and female reproductive tissues and cell types as the events of meiosis and gametophyte formation progress through fertilisation and early post-fertilisation events of seed initiation.

In cowpea, as in other angiosperms, the male and female gametes develop in a defined temporal sequence within the developing male anther and female ovule in the flower (Salinas-Gamboa et al., 2016). Gametogenesis involves meiotic reduction, mitosis and cell differentiation to produce the multicellular pollen grain in the anther and female gametophyte in each ovule. During double fertilisation, sperm cells individually fuse with the egg cell and fused polar nuclei in the central of the female gametophyte giving rise to the embryo and nutritive endosperm, respectively. The endosperm is utilised during embryo formation and seed storage protein reserves are stored in legume cotyledons.

Here, we report on a novel cowpea gene expression platform, VigExp, that incorporates transcriptomes of major vegetative tissues from IT97K-499-35, including a developmental time series for pods and seeds (VuGEA; Yao et al., 2016) and dissected floral tissues from a developmental time series during male and female gametogenesis (Spriggs et al., 2018). Novel, laser-captured cell types, collected in a developmental series spanning the events of male and female gamete formation and early embryogenesis, not previously reported for any legume species, are also incorporated. VigExp proved reliable and robust for establishing and visualising cowpea gene expression patterns following verification using RT-qPCR, *in situ* localisation, and whole plant transformation. Cowpea homologs of agronomically important legume nodulation genes exhibited similar cell/tissue-type expression patterns, with *VuNFR5*, a critical receptor for rhizobia Nod factor perception, expressed in a wider range of tissues than reported for other legumes. The cowpea homologue of the parthenogenic gene, *BABYBOOM-like* (*BBML*), also exhibited an expression pattern consistent with its functionality. Moreover, knockout of the cowpea homolog of the meiosis-associated gene, *TARDY ASYNCHRONOUS MEIOSIS* (*TAM*), revealed a critical role for this A-type cyclin in male meiosis. Collectively, these findings further validate the robustness of the platform. Moreover, utility of VigExp is demonstrated through identification of novel reference candidates for gene expression analyses, constitutively expressed genes providing promoters for genetic transformation, and tissue- and cell-specific expressed genes providing promoters for targeting these regions. VigExp represents a robust platform for supporting the identification and testing of genes that aid in the improvement of cowpea and related legume crops. Given the inclusion of additional developmentally staged reproductive cell types and tissues, VigExp can help identify relevant candidate genes acting in several developmental pathways of interest, including those that alter cowpea reproduction from a sexual to an asexual mode of seed formation (synthetic apomixis).

## RESULTS AND DISCUSSION

### Mapping of raw transcriptome data from whole cowpea tissues

Forty-three previously generated RNA sequencing datasets from vegetative and reproductive tissues (illustrated in Figure 1A and Supplemental Figures 2A and 3) collected from the cowpea variety IT97K-499-35 by Yao et al., (2016) and Spriggs et al., (2018) were remapped to the IT97K-499-35 reference genome (Lonardi et al., 2019). The total number of uniquely mapped reads within each sample ranged from 3,920,208 to 36,041,470, and 20,969,820 reads on average (Supplemental Table 1). Sample normalisation achieved using Gene length corrected trimmed mean of M-values (GeTMM) revealed high-quality normalisation across the samples (Supplemental Figure 2A). T-distributed stochastic neighbour embedding (t-SNE) established the replicate samples clustered closely amongst the respective tissue types (Figure 1B), demonstrating robust GeTMM normalisation and RNAseq datasets.

**Figure 1.**
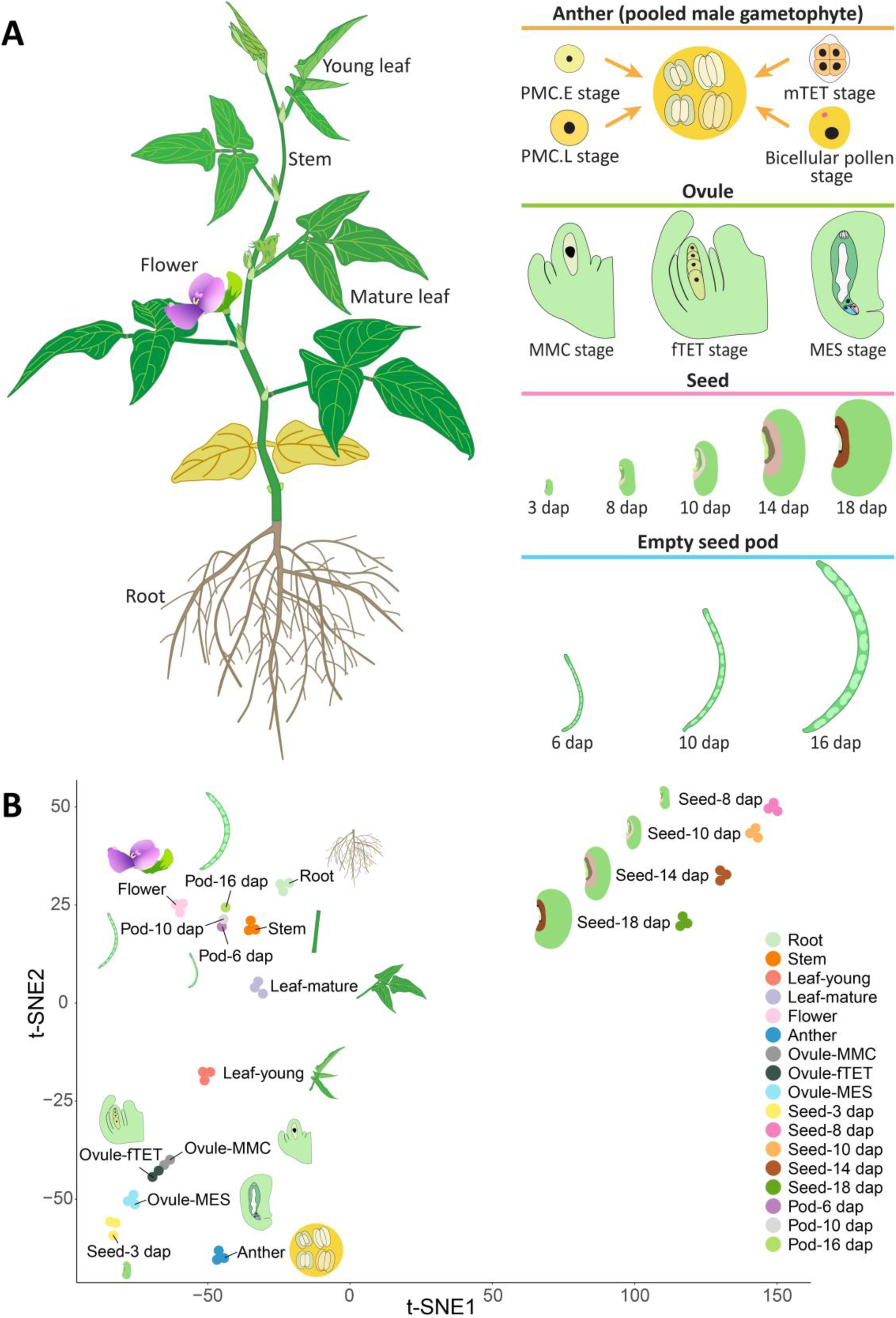
Cowpea transcriptomes from various tissue types and stages verified by sample clustering. **(A)** Cowpea plant and tissues with text illustrating various tissues collected (Supplemental Figure 3). **(B)** t-SNE plot of transcriptomes from different tissues/stages. Replicate number per stage ranged from one to three (Supplemental Figure 3, Whole tissue samples), with dots of the same colour representing biological replicates. Abbreviations: PMC.E: pollen mother cells at early stage; PMC.L: pollen mother cells at later stage; mTET: male tetrads; MMC: megaspore mother cell; fTET: female tetrads; MES: mature embryo sac at anthesis; dap: days after pollination.

### Generation of cowpea reproductive cell-type specific transcriptomes

The cowpea variety IT86D-1010 was used to generate 13 different reproductive cell-type transcriptomes (two biological replicates) from staged male and female reproductive tissues, and early embryogenesis stages, collected using laser capture microdissection (LCM), or in the case of sperm cells, via tissue collection and centrifugation (Figure 2A; Supplemental Figure 3; Gursanscky et al., 2020). The datasets were normalised and mapped to the IT86D-1010 genome (https://data.csiro.au/collection/csiro:63431), which has 93.75% conservation with the coding sequences of IT97K-499-35 (Phytozome annotated *V. unguiculata* genome v1.2).

**Figure 2.**
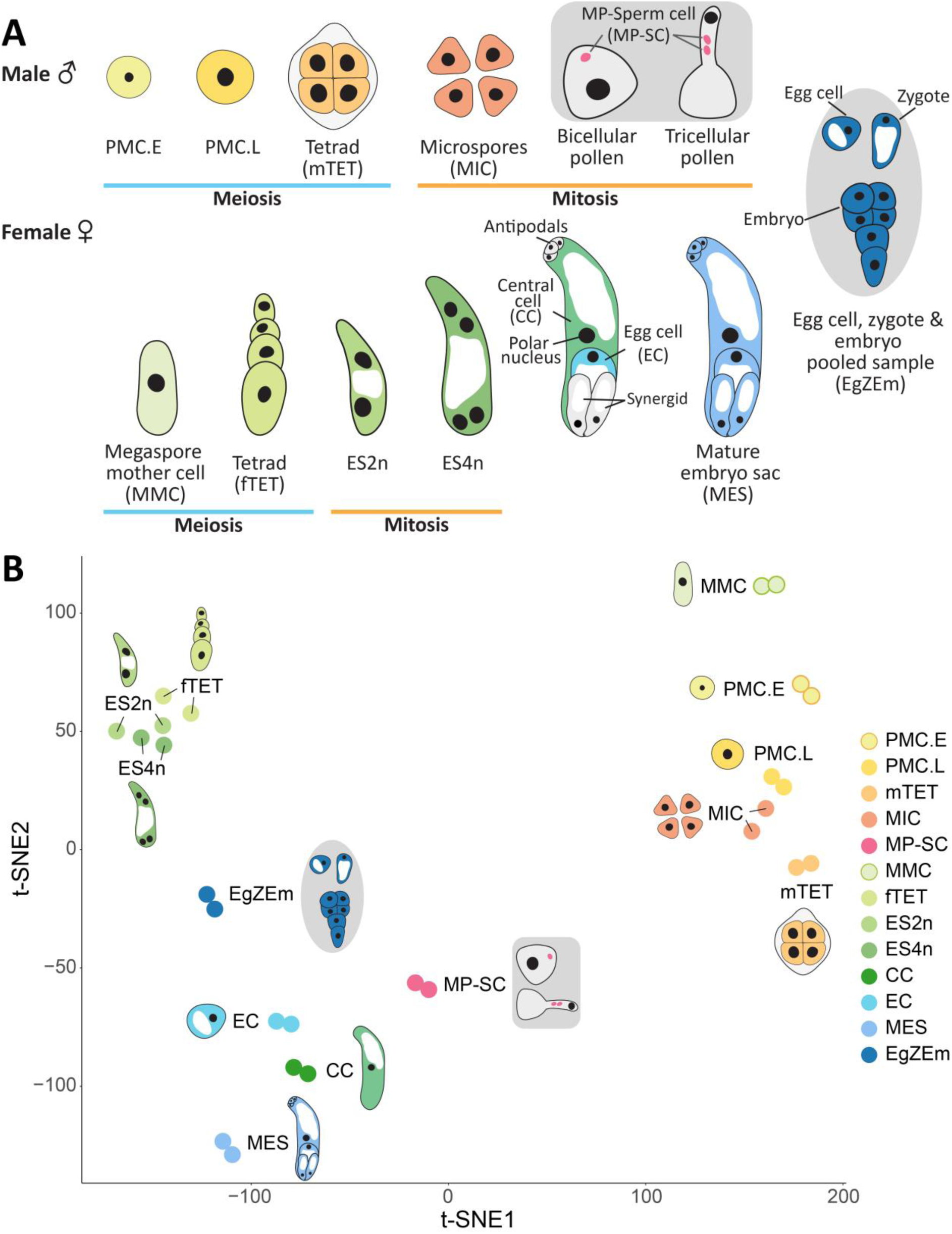
Cowpea transcriptomes from various stages of male and female reproductive development verified by sample clustering. **(A)** Male and female gametophyte development in cowpea showing cell-type stages collected by laser capture microdissection (LCM) or mature pollen-sperm cell sampling (Supplemental Figure 3). **(B)** t-SNE plot of transcriptomes from male and female cell types (*n* = 2), with dots of the same colour representing biological replicates. Grey shading represents samples having multiple developmental stages. Abbreviations: PMC.E: pollen mother cells at early stage; PMC.L: pollen mother cells at later stage; mTET: male tetrads; fTET: female tetrads; ES2n: embryo sacs with two nuclei; ES4n: embryo sacs with four nuclei; MP-SC: mature pollen-sperm cell.

Male gametophyte transcriptomes obtained from pollen mother cells include early and late stages of meiosis I (PMC.E, PMC.L), microspore tetrads (mTET), and uninucleate microspores (MIC) (Figure 2A). In mature anthers, only 9.7% of mature pollen grains contained two sperm cells (*n* = 500). This indicates final generative cell mitosis giving rise to two sperm cells primarily occurs following pollen tube germination, a phenomenon commonly observed in legumes and 70% of flowering plant species overall (Brewbaker, 1967; Haerizadeh et al., 2009; Williams et al., 2014). Therefore, anthers and pistils were harvested at anthesis to create a mixed sperm cell-enriched sample via lysis and centrifugation. This sample is referred to here as the mature pollen-sperm cell sample (MP-SC; Figure 2A Male, shaded in a rounded grey rectangle).

Female gametophyte-related cell-type and early embryo samples included the megaspore mother cell (MMC) undergoing meiosis, the subsequent tetrad of post-meiotic megaspores (fTET), and developing mitotic embryo sacs having two or four nuclei (ES2n, ES4n) (Figure 2A). The mature embryo sac (MES) containing the egg cell (EC), the flanking synergids (Sy) and fused central cell (CC) nuclei prior to anther dehiscence, at the time of stamen emergence. Individual EC, CC and mature embryo sac (MES) samples were also collected (Figure 2A, Female). The early embryo (Em) sample was a mixed sample (EgZEm) comprising tissue from mature eggs, developing zygotes and early globular stages of cowpea embryogenesis (Figure 2A, shaded in a grey oval).

RNAseq reads generated from the reproductive cell types were quantified from 31,948 genes predicted by Phytozome and normalised following transcript per million reads (TPM) conversion and GeTMM normalisation (Supplemental Figure 2B). The total number of uniquely mapped reads within each sample ranged from 7,430,663 to 23,443,429, and 15,010,763 reads on average (Supplemental Table 1). The t-SNE analyses confirmed the sample quality and following normalisation the biological replicates were found to cluster, further demonstrating the datasets were of high quality (Figure 2B).

### Validation of transcriptomic datasets

Validation of the transcriptomic datasets was achieved using RT-qPCR, with cDNA isolated from whole tissues collected at comparable developmental stages to the whole, LCM and MP-SC samples used to generate the transcriptomes (Figures 1, 2; Supplemental Figure 3; Supplemental Table 1). To normalise the RT-qPCR results for each candidate gene investigated, reliable reference genes first needed to be identified. Eight candidate reference genes were selected, including seven that exhibited consistent expression profiles throughout the transcriptomic datasets, *VuCAM6* (*Calmodulin 6*), *VuUBA1* (*Ubiquitin-activating enzyme 1*), *VuFBX2* (*F-box protein 2*), *VuSWAP70* (*Switch-Associated Protein 70*), *VuAAC3* (*ADP/ATP carrier 3*), *VuAGB1* (*GTP binding protein beta 1*), and *VuCPA* (*Capping protein A*), and one previously reported in the literature, *VuADF* (*Actin depolymerizing factor 4*; Juranić et al. 2020). Expression of these candidate genes was assessed via RT-qPCR using quantified mRNA isolated from whole tissue samples across 26 tissue types. The Cq/gene expression results were then subjected to several analyses (outlined in the Methods section) to establish the most robust for use as reference genes (Supplemental Figure 4 and Figure 3A), with *VuUBA1* and *VuFBX2* paired genes identified as most reliable for use as reference genes to validate the transcriptomic datasets (Figure 3A right).

**Figure 3.**
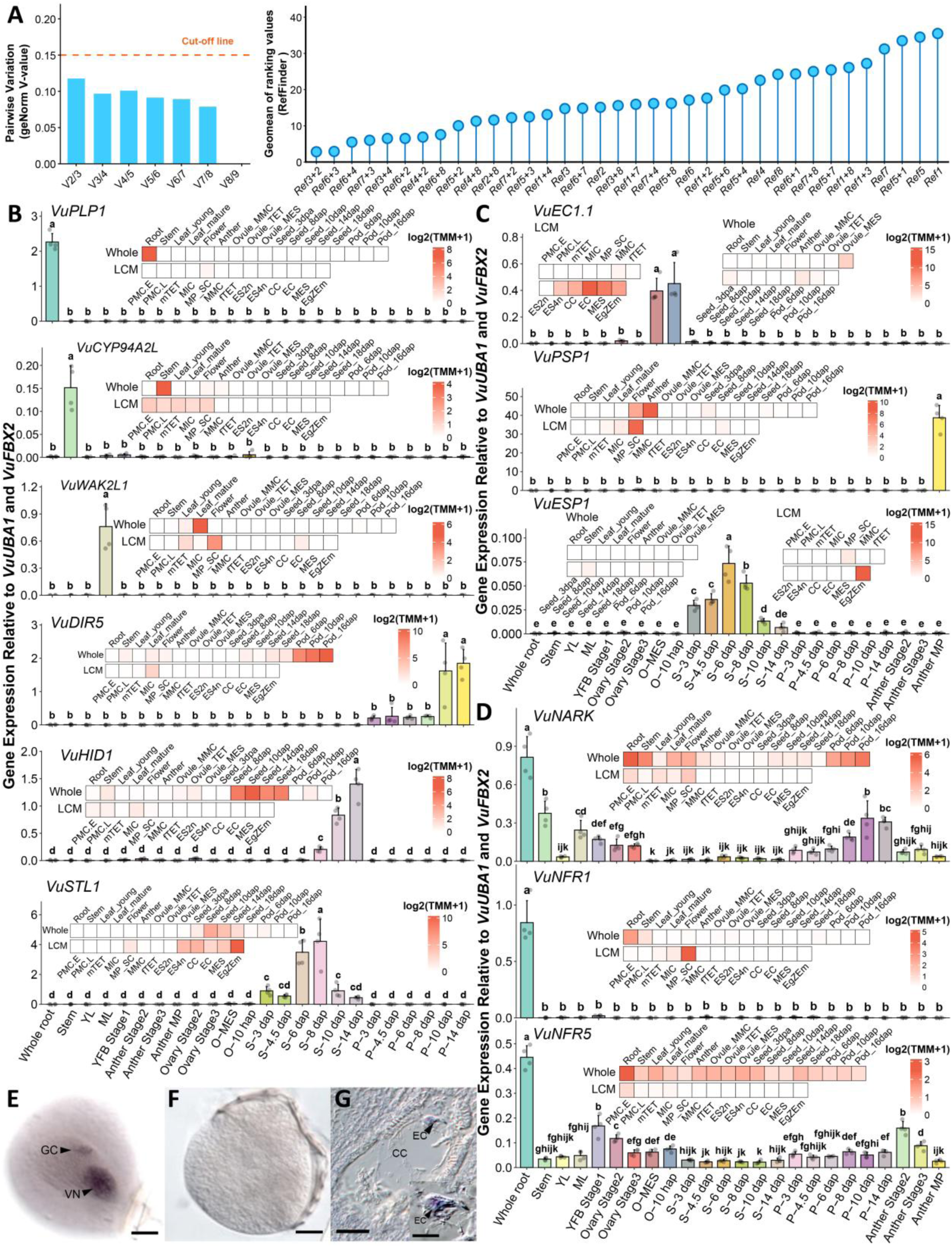
Identifying reference genes of cowpea, and validation of cowpea transcriptomes through RT-qPCR and *in situ* hybridisation of selected genes. (A) Gene pairwise variation (Vn/Vn+1) (left) to determine the number of reference genes for accurate Cq normalisation and gene stability ranking (right) of single/paired candidate reference genes. Transcript abundance determined via RT-qPCR of candidate genes identified in **(B)** whole tissues and **(C)** reproductive cell types, and of **(D)** cowpea homologs of known legume nodulation genes from IT86D-1010. The *in situ* hybridisation of (E) *VuXTH6* mRNA in mature pollen generative cell (the precursor of two sperm cells) and vegetative nucleus, **(F)** with no signal in the sense control, and (G) *VuEC1.1* mRNA in a mature embryo sac. Ref1: *VuCAM6*; Ref2: *VuUBA1*; Ref3: *VuFBX2*; Ref4: *VuSWAP70*; Ref5: *VuAAC3*, Ref6: *VuAGB1*; Ref7: *VuCPA*; Ref8: *VuADF*. Abbreviations: GC: generative cell; VN: vegetative nucleus; EC: egg cell; CC: central cell; PMC.E: pollen mother cells at early stage; PMC.L: pollen mother cells at later stage; mTET: male tetrads; fTET: female tetrads; ES2n: embryo sacs with two nuclei; ES4n: embryo sacs with four nuclei; MP-SC: mature pollen-sperm cell; YL: young leaf; ML: mature leaf; YFB: young flower bud; MP: mature pollen; O: ovule; MES: mature embryo sac; hap: hour after pollination; S: seed; P: pod; dap: day after pollination. Scale bars: **E - F** = 10 µm, **G** = 18 µm (inset = 10 µm).

Six candidate genes, *VuPLP1* (*Patatin-related phospholipase A*), *VuCYP94A2L* (*Cytochrome P450 94A2-like*), *VuWAK2L1* (*Wall-associated kinase 2-like 1*), *VuDIR5* (*Dirigent protein 5*), *VuHID1* (*2-hydroxyisoflavanone dehydratase 1*), and *VuSTL1* (*Scorpion toxin-like 1*), exhibiting tissue-specific expression were selected from the whole tissue transcriptomes (Figure 3B heatmaps). All six were confirmed to be highly specific in expression following RT-qPCR (Figure 3B barplots), demonstrating the robust quality of the transcriptomic data. An additional three candidate genes, *VuEC1.1* (*Egg cell-secreted protein 1.1*), *VuPSP1* (*Plant self-incompatibility protein S1*), and *VuESP1* (*Embryo secreted protein 1*), exhibiting cell-type-specific expression were selected from the LCM and MP-SC transcriptomes (Figure 3C heatmaps). Many of these candidates were also highly specific in their expression following RT-qPCR, with some exhibiting very low levels of expression (Figure 3C barplots). This highlights the quality of the LCM and MP-SC transcriptomes. It also underscores some of the cell-specific expression of the candidate genes, which were diluted in the mixed cell types of whole tissues collected for RT-qPCR, compared with the individual cell types obtained by LCM and MP-SC for the transcriptomes. Expression levels of additional six genes, *VuEXT2L1* (*Extensin-2-like 1*), *VuHRD1* (*Hardy, AP2_ERF domain-containing protein 1*), *VuEC1.2* (*Egg cell-secreted protein 1.2*), *VuEC1.3* (*Egg cell-secreted protein 1.3*), *VuRHS14* (*Pectate lyase 1/Root hair specific 14*), and *VuXTH6* (*Xyloglucan endotransglucosylase/hydrolase 6*), identified from whole tissues and cell-types further confirmed the robust quality of the transcriptomic datasets (Supplementary Figure 5).

Well-known gene candidates from the legume-rhizobia symbiosis were also investigated. This includes the Nod factor receptor genes, *NFR1* and *NFR5*, that enable the host plant to recognise their compatible rhizobia partner, and the autoregulation of nodulation receptor, *NARK*, which enables the host to control how many nodule structures it forms (Ferguson et al., 2019). *NARK*, which is reported to be expressed in roots and leaves of several legume species (including soybean, common bean, pea, *Medicago truncatula*, and *Lotus japonicus*), was also found to be widely expressed in roots and leaves of cowpea (Figure 3D). Interestingly, it was also expressed in stem, ovary and pod tissues, suggesting *VuNARK* may function more broadly throughout the plant than previously thought. *NFR1* and *NFR5* are reported to be specifically expressed in root tissues, where they perceive the rhizobia-produced Nod factor signal. Here, we found *VuNFR1* exhibits this root-specific expression pattern, but *VuNFR5* was observed in both transcriptomic and RT-qPCR samples to be expressed throughout the plant (Figure 3D). This suggests additional roles for *VuNFR5* in various tissues, likely in complexes with factors other than *VuNFR1*.

Further validation was achieved by assessing the location of candidate gene mRNA transcripts in mature pollen-sperm cell samples and developing ovule cell types using *in situ* hybridisation. *VuXTH6* mRNA was detected at high levels in the generative cell and vegetative nucleus, with low levels detected in the vegetative cell cytoplasm (Figure 3E). Whereas transcripts from female gametophyte-specific *VuEC1.1* were highly enriched in the egg cell, but not the central cell (Figure 3G). Collectively, these findings provided confidence in the integrity of the isolated reproductive cell type transcriptomes.

### Development of VigExp to visualise transcriptome datasets of cowpea

A user-friendly platform for integrating and viewing cowpea transcriptome data with no programming requirements was developed, called VigExp (Figure 4). This was achieved according to a designed bioinformatic workflow (Supplemental Figure 6).

**Figure 4.**
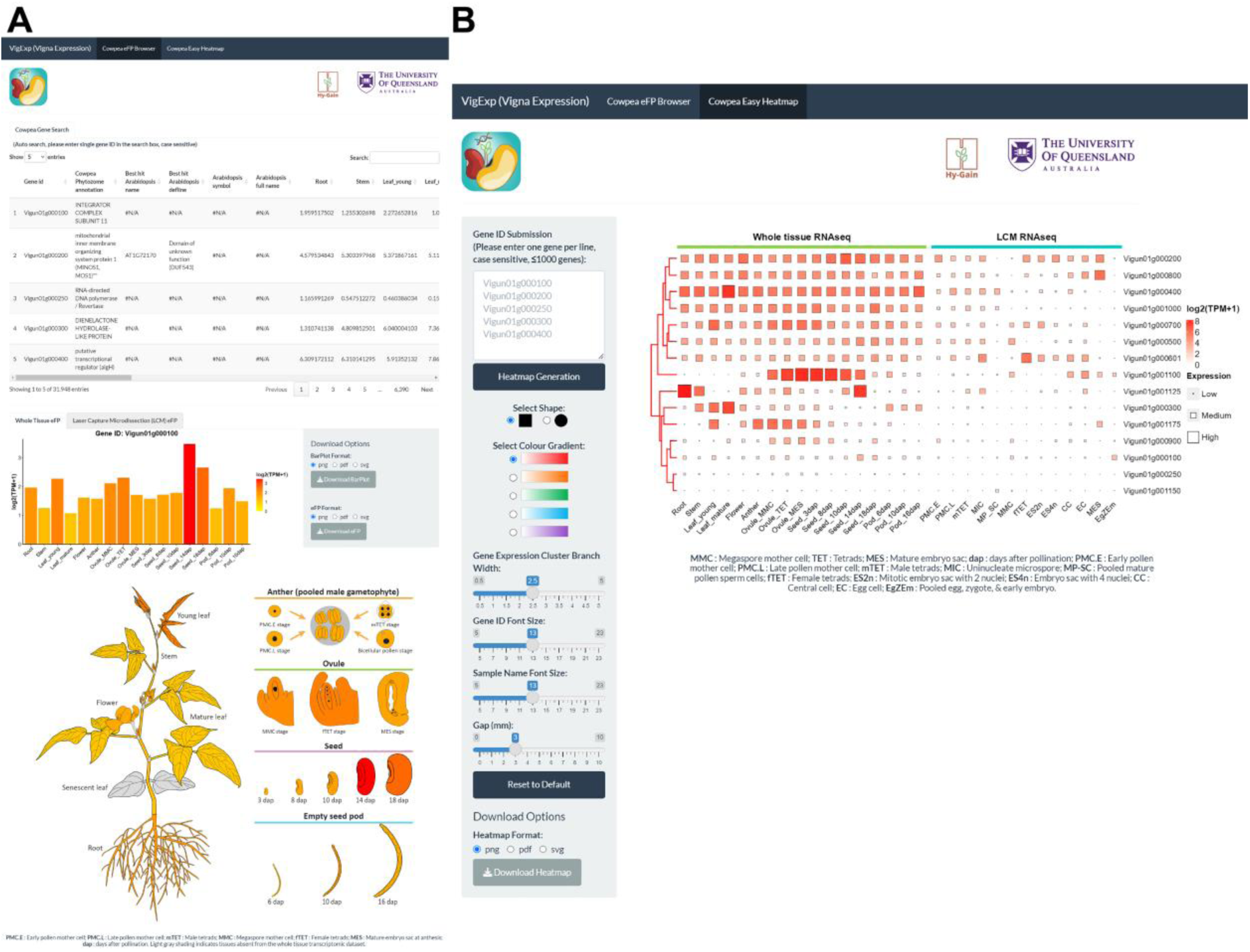
VigExp interface to search and visualise gene expression profiles. **(A)** The cowpea gene search and eFP Browser function in VigExp. **(B)** The ‘Cowpea Easy Heatmap’ function in VigExp.

Due to differences in the sample collection method and sample type (i.e. whole tissue or LCM single cell), batch effects were likely to be introduced that could not be eliminated by normalisation. Therefore, the VigExp tool hosts separate normalisation matrices for whole tissue datasets and LCM datasets. The whole tissue matrix includes 43 log2 (TPM+1) datasets and the LCM matrix includes 26 datasets (Supplemental Figure 3). In total, 31,948 annotated genes from the IT97K-499-35 in Phytozome (https://phytozome-next.jgi.doe.gov/) can be viewed in VigExp. In addition, 18,513 genes from IT97K-499-35 have a potential gene ortholog identified in *Arabidopsis*, which can also be viewed in VigExp along with the associated gene annotation and functional description from TAIR (https://www.arabidopsis.org/). This information can be searched using the gene ID, name, or annotation in the interactive table in VigExp (Figure 4). Moreover, the TPM matrix can be copied or downloaded for further downstream analyses, such as generating heatmaps. Bar plots and Whole Tissue eFP Browser/LCM eFP Browser (Figure 4A) enable the expression profile of genes to be visualised in histogram or tissue diagrams. The ‘Cowpea Easy Heatmap’ function (Figure 4B) also enables fast visualisation of many gene expression profiles simultaneously (up to 1,000 cowpea gene IDs), ideal for visualising gene families, tissue- or cell-specific genes, and more. Pre-designed settings to customise the heatmaps generated are also available. High-quality “png (600 dpi)”, “svg”, or “pdf” graphs can be generated and downloaded for both the cowpea eFP Browser and the Easy Heatmap (Figure 4). Moreover, VigExp can be adapted and updated to incorporate additional transcriptome datasets in future.

### Validation of VigExp functionality

To validate the functionality of VigExp and confirm its utility, expression of the cowpea orthologue of *BABYBOOM-like* (*BBML*) was identified. Members of the *BBML* gene family can induce parthenogenesis in monocots, such as rice and sorghum when ectopically expressed in the egg cell (Khanday et al., 2019; Simons et al., 2025). These genes are typically expressed during pollen development, with fertilisation introducing the gene product into the egg to aid embryo initiation. Here, we utilised the VigExp platform to assess expression patterns of the cowpea *VuBBML* gene family using the ‘Cowpea Easy Heatmap’ feature (Supplemental Figure 7) and identified the cowpea *BBML* homolog (*VuBBML1*), which is expressed during pollen development and post-fertilisation during early embryogenesis (Figure 5). This pattern strongly correlates with *BBML* expression reported for other species and is consistent with its functional role in parthenogenesis when directed to the egg cell. RT-qPCR validation also confirmed the consistency of *VuBBML1* expression identified in the transcriptome datasets, being predominantly in the root, young flower buds (YFB Stage 1), and 3 - 14 dap seeds.

**Figure 5.**
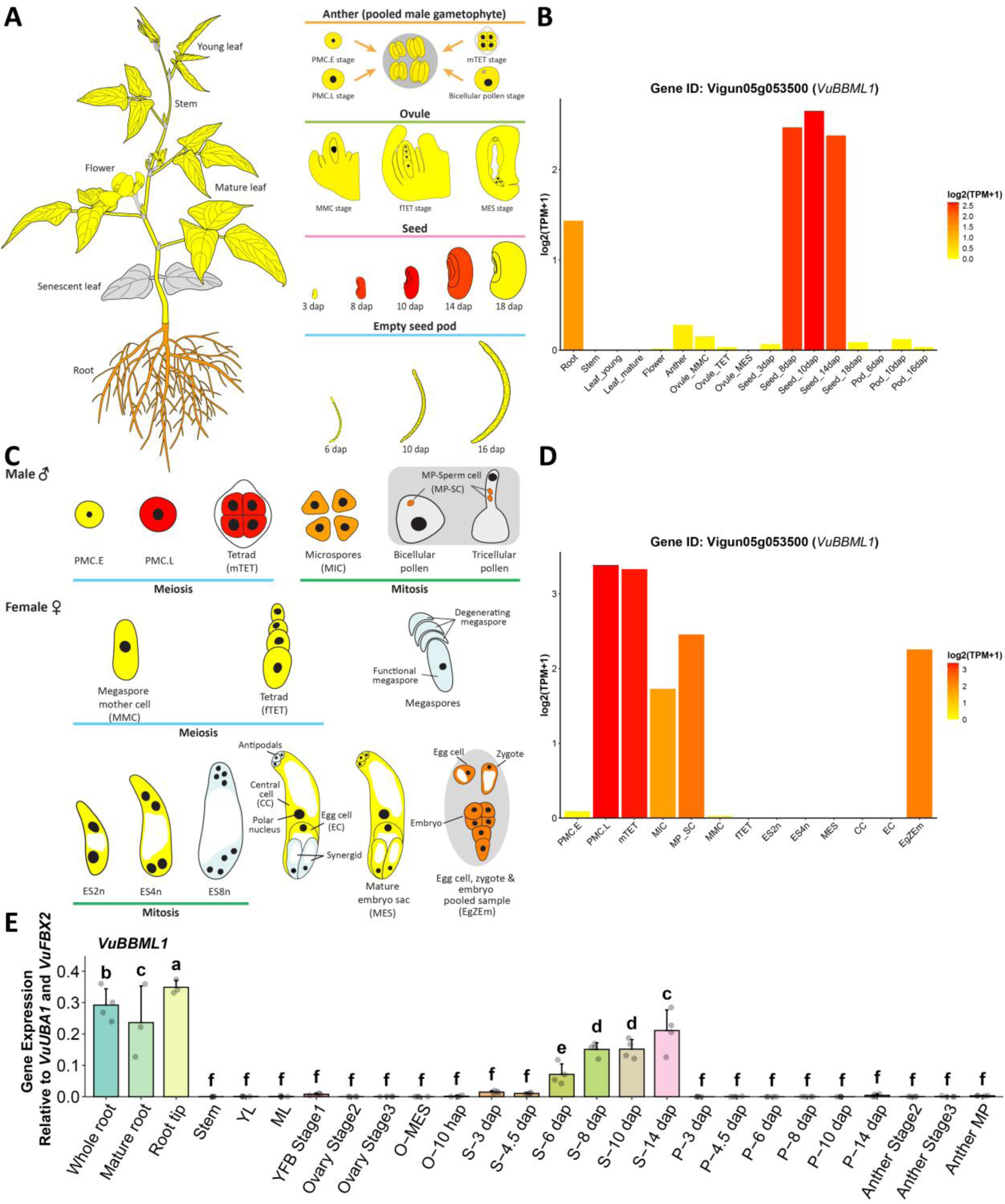
Expression pattern of the cowpea homolog of *BBML* visualised using the VigExp platform. **(A)** Location of expression in whole tissues, including **(B)** Expression levels in these tissues. **(C)** Location of expression in reproductive cell types, including **(D)** Expression levels in these cells. **(E)** RT-qPCR expression of *VuBBML1* in various whole tissues from IT86D-1010.

To further demonstrate the utility of VigExp, the platform was used to help identify genes exhibiting constitutive expression patterns. Promoters from these genes can be useful for establishing the efficiency of transformation and avoiding chimeric sectors when conducting genetic modification and gene editing studies. Genes exhibiting high expression at the cotyledonary node and in subsequent developing shoots were targeted here. Three candidate genes, *pVuEF1a*, *pVuUBQ4* and *pVuUBQ10*, were identified (Figure 6A). The promoter of *VuEF1a* was very strongly expressed throughout the plant and used here to drive spectinomycin resistance in vectors for genetic transformation (Figure 6B-C). Promoters from *VuUBQ4* and *VuUBQ10* were fused to *ZsGreen* and their expression examined in transgenic cowpea. The promoter of *VuUBQ4* was identified as an ideal candidate for transgenic studies due to its strong ability to drive *ZsGreen* expression in the shoot apical meristem, developing shoots, mature leaves, trichomes, roots, flower buds and pollen (Figure 6D-J; Supplementary Figure 8). The promoter of *VuUBQ10* also provided high expression throughout the plant (Figure 6K-P; Supplementary Figure 9). Interestingly, *VuUBQ4* was highly expressed in the hilum, whereas *VuUBQ10* was highly expressed in the seed cotyledon (Figure 6Q-V, P), allowing quick selection of dry transgenic seeds. Collectively, these studies demonstrate the utility of VigExp in helping identify gene and promoter candidates of interest for facilitating cowpea crop improvement through genetic transformation.

**Figure 6.**
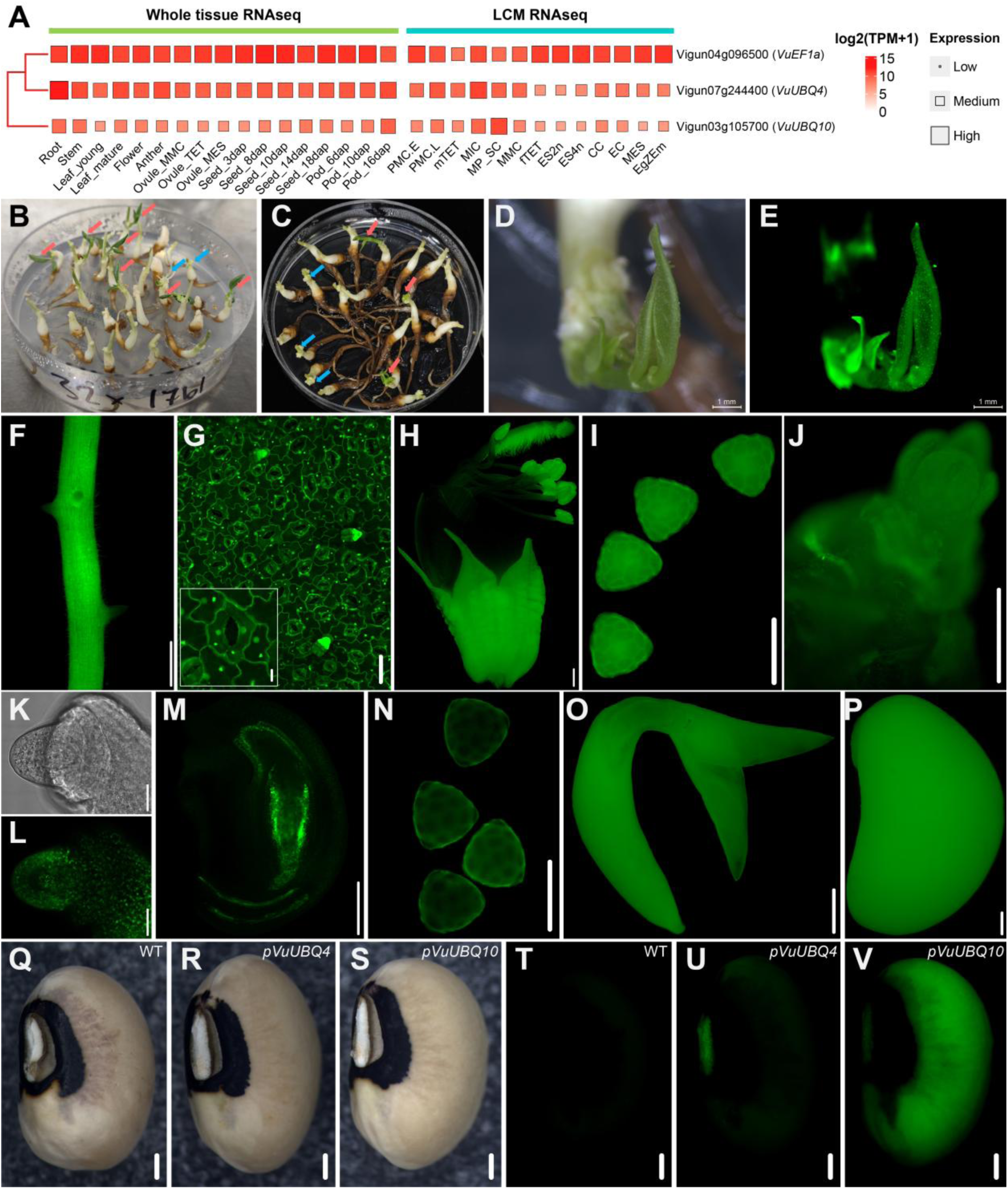
Cowpea constitutive promoter identification and utility in cowpea transformation. **(A)** Heatmap generated by VigExp shows three highly expressed candidate genes whose promoters were used here for developing cowpea-specific transformation vectors. **(B and C)** *pVuEF1a:: CTP-SpcN:: tVuEF1a* and *pVuEF1a:: CTP-aadA1:: tVuEF1a* used in transgenic cowpea selection on spectinomycin shoot induction media (red arrow: transformed dark green shoots, blue arrow: non-transformed bleached shoots). **(D - J)** *pVuUBQ4*:: *ZsGreen:: tVuUBQ4* used as a visualisation marker in **(D)** young shoots in bright field and **(E)** dark field, and in **(F)** root, **(G)** mature leaf and trichomes, **(H)** dissected floral bud at stage 1F-IX, **(I)** mature pollen**, (J)** shoot apical meristem**; (K - P)** *pVuUBQ10*:: *ZsGreen:: tVuUBQ10* expression in **(K)** the ovule at megaspore mother cell in bright field and **(L)** dark field, and in **(M)** mature embryo sac stage, **(N)** mature pollen**, (O)** embryonic axis, and **(P)** cotyledon. **(Q - V)** Seed of IT86D-1010 **(Q, T)** wild-type, **(R, U)** *pVuUBQ4*:: *ZsGreen:: tVuUBQ4* transformed and **(S, V)** *pVuUBQ10*:: *ZsGreen:: tVuUBQ10* transformed in **(Q-S)** bright field and **(T - V)** dark field. Wild-type IT86D-1010 was used as negative control to adjust microscope settings to avoid detection of autofluorescence. Scale Bars: **D - H, O - V** = 1000 μm; **I, M - N** = 100 μm; **J** = 200 μm; **K - L** = 20 μm.

### CRISPR editing of *VuTAM2*, identified using VigExp, results in the omission of meiosis II during microsporogenesis

VigExp was also used to develop a cowpea CRISPR/Cas9 gene editing system. To validate this work, the homolog of *Arabidopsis AtTAM* and tomato *SlTAM*, called *VuTAM2*, was identified in cowpea. A phylogenetic tree (Supplemental Figure 10) and heatmap of the TAM protein family generated using VigExp (Supplemental Figure 11), identified high expression of *VuTAM2* in both the PMC.L and MMC (Figure 7A), indicating potential function during meiosis. Gene knockout using CRISPR was conducted using IT86D-1010 to further evaluate *VuTAM2* function during meiosis in cowpea. Three *VuTAM2* biallelic CRISPR knockout T1 plants were identified from five T0 quality events: G23-T1_plant_#4-S1, G23-T1_plant_#30-S14, G23-T1_plant_#30-S15 (Figure 7J - L; Supplemental Figure 12). Pollen from these plants exhibited 100% dyad formation from most flowers, with only a few flowers from each line producing a mixture of tetrads and dyads (Figure 7C - D). However, these *VuTAM2* knockout lines develop tetrad ovules, indicating meiosis occurs normally in female gametogenesis (Figure 7E - F). Pod formation was also greatly reduced in the edited lines (Figure 7J - L; Supplemental Table 3), but seed setting per pod was not affected (Figure 7M; Supplemental Table 3). Mature pollen exhibited a considerable amount of non-viable pollen grains (Figure 7G - H), with viable pollen from single anthers ranging from 0 to 35.4% (Figure 7I). Similar issues with reduced pollen viability were also reported for tomato *SlTAM* CRISPR knockout lines (Wang et al., 2024).

**Figure 7.**
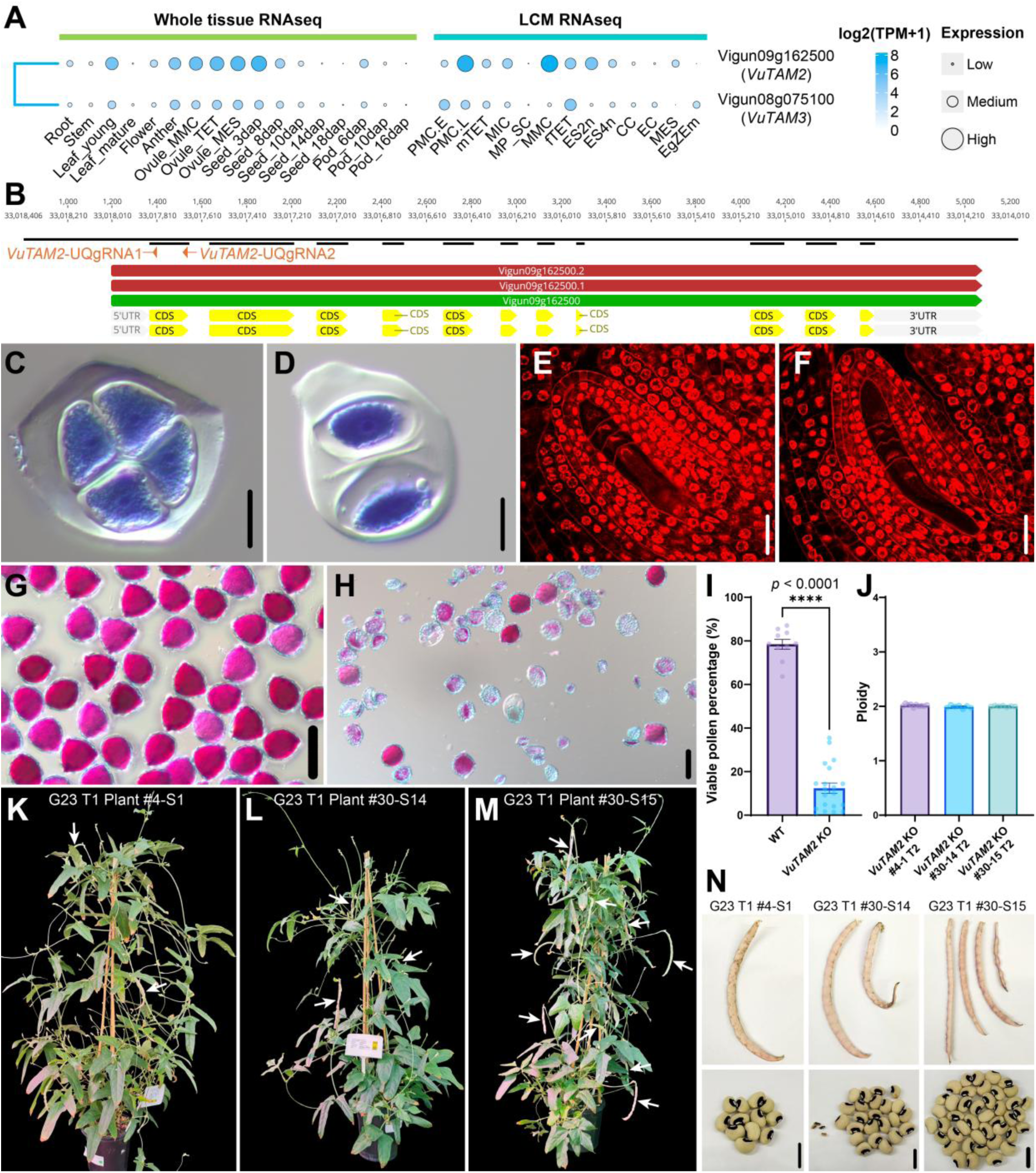
Cowpea *VuTAM2* CRISPR editing results in the omission of meiosis II during microsporogenesis. **(A)** *VuTAM2* and *VuTAM3* expression heatmap profiles generated by VigExp. **(B)** Two gRNAs were designed to target the first exon in the *VuTAM2* coding sequence. **(C)** Tetrad pollen in the wild-type IT86D-1010. **(D)** Microsporocyte dyads instead of tetrads in *VuTAM2* biallelic CRISPR knockout lines. **(E)** Female tetrad in wild type IT86D-1010. **(F)** Female tetrad also observed in *VuTAM2* biallelic CRISPR knockout lines. **(G - H)** Pollen viability determined using Alexander’s staining in **(G)** wild type and **(H) a** *VuTAM2* biallelic knockout line, where bright pink represents viability. **(I)** Male fertility in wild-type and *VuTAM2* biallelic knockout lines. **(J)** Ploidy determination in *VuTAM2* knockout T2 progenies. **(K - M)** Three *VuTAM2* biallelic knockout lines showing extensive sterility, with only a few pods forming (white arrows). **(N)** Mature pods and seeds from *VuTAM2* biallelic knockout lines. Scale Bars: **C - F** = 20 μm; **G - H** = 100 μm; **N** = 1000 μm.

To determine the ploidy of *VuTAM2* knockout lines, a novel method was developed for establishing cowpea ploidy based on flow cytometry with internal standardisation (Supplemental Figure 13). Reference genomes from five different species, including common bean (*Phaseolus vulgaris*), sorghum (*Sorghum bicolor* Tx430), soybean (*Glycine max* var. Bunya), popping maize (*Zea mays* var. Everta), and sunflower (*Helianthus annuus*), were tested. Comparison of cowpea samples with the reference genome samples enabled an internal correction for normalisation. To confirm the method is robust, haploid, diploid and tetraploid Tx430 sorghum were used for verification, as there are currently no polyploid cowpea lines available. Findings established that the best reference genomes for normalising ploidy were soybean for cowpea and popping maize for sorghum, as peaks for the other species tested either overlapped or were too far apart to be suitable as reference genomes (Supplemental Figure 13). Three to four flow cytometry peaks were usually observed, and only Mean-x values of the G_0_/G_1_ peak were used for cowpea or sorghum downstream normalisation (Supplemental Figure 13A and B). Haploid, diploid and tetraploid Tx430 sorghum ploidy verification was significantly improved following normalisation with the reference species (Supplemental Figure 14A and B). Consistent results were also achieved with diploid cowpea (Supplemental Figure 14C and D), further demonstrating the robustness of the method.

Following verification of the technique, the ploidy level of *VuTAM2* knockout lines was determined. Flow cytometry confirmed T2 seedlings (seven - ten for each line) from all three events were diploid (Figure 7N, Supplemental Table 6), indicating the progeny were most likely derived from the few flowers that produced some tetrad (n) pollen fertilising the female tetrads (n). Collectively, these results demonstrate an important role for *VuTAM2* in pollen meiosis, with no, or possibly a redundant, role in female gametogenesis.

### Concluding remarks

In this study, we developed a comprehensive tool for visualising cowpea transcriptomic data, VigExp. Transcriptomes of reproductive cell types that are challenging to collect are reported for the first time in any plant species. Together with additional transcriptomes generated from whole tissues, their inclusion in the VigExp platform enabled the identification of new reference genes, constitutive promoters, several cell/tissue-type specific promoters, meiosis-related genes, and novel developmental gene candidates, which represent new enabling resources for the scientific community. Using this tool, a cowpea transformation vector system for transgene insertion or CRISPR gene editing was established. To validate this system, *VuTAM2* was functionally characterised by generating stable knockout mutants, revealing an essential role for the A-type cyclin in cowpea male, but not female, meiosis, with no, or possibly a redundant, role in female gametogenesis of cowpea. Importantly, VigExp and the associated cowpea eFP browser and heatmap are designed to be adaptable, enabling additional transcriptomic datasets to be added as they become available to maintain the relevance of these bioinformatic tools in future. Moreover, due to the high genomic similarity in the *Fabaceae* family (Sprent, 2007; Supplemental Figure 1), particularly amongst major legume crops (e.g. cowpea, soybean, common bean, pea, lentil, chickpea, faba bean, mung bean), findings from one species can often be extrapolated to other species based on genomic microsynteny and gene sequence similarity (e.g. Hastwell et al., 2017; Zhang et al., 2021). Hence, findings using the VigExp and eFP browser/Easy Heatmap tools can benefit cowpea directly and legume crops more broadly. Collectively, VigExp and the novel genetic factors identified here can help underpin future strategies that interrogate and functionally characterise genes and pathways in economically significant legume crops.

## METHODS

### Raw transcriptome data sourced for whole cowpea tissues

Raw transcriptome datasets from whole tissues of cowpea variety IT97K-499-35 were obtained from NCBI: PRJNA389300 (Yao et al., 2016), including root, stem, mature leaf, flowers at anthesis, empty seed pods collected 6, 10 and 16 days after pollination (dap), and staged seeds collected 8, 10, 14 and 18 dap. Additional datasets were obtained from the CSIRO Data Access Portal (https://data.csiro.au/collection/csiro:27153v3; Spriggs et al., 2018), including young unexpanded leaves plus whole reproductive tissues, including pooled anthers containing a variety of male gametophyte developmental stages, staged ovules containing pooled male gametophyte (Anther-PMG), megaspore mother cells (Ovule-MMC), female meiotic tetrads (Ovule-fTET), mature female gametophytes (Ovule-MES), and staged seeds collected 3 dap.

### Plant growth and reproductive cell collection

Plants of cowpea variety IT86D-1010 were grown under previously described glasshouse conditions (Salinas-Gamboa et al., 2016). LCM was used to isolate individual cell types to avoid limitations of other single-cell methodologies (Haghan et al., 2026), such as incompatibility of large reproductive cell types with smaller 10 × flow cell channel (Washburn et al., 2023) and the very limited cell-type and developmental staging markers in legumes, which can lead to inaccurate identification of cell clusters and developmental stages in downstream single-cell/nucleus RNAseq analysis. Sample preparation was performed using published protocols (Rasheed et al., 2006; Tucker et al., 2012; Okada et al., 2013) modified for isolating gametogenic cell types of cowpea. In brief, flowers at different developmental stages were fixed in 3: 1 ethanol: acetic acid and embedded in butylmethyl-methacrylate. Sections (5 µm) were placed on membrane slides (Leica), treated with acetone and dissected using an AS-LMD laser microscope (Leica; Tucker et al., 2012). Thirteen specific cell-types were collected, including megaspore mother cell (MMC), female tetrads (fTET), mitotic embryo sac with 2 nuclei (ES2n), embryo sac with 4 nuclei (ES4n), central cell (CC), egg cell (EC), mature embryo sac (MES), pooled egg, zygote, and early embryo (EgZEm), early pollen mother cell (PMC.E), late pollen mother cell (PMC.L), male tetrads (mTET), uninucleate microspore (MIC), and mature pollen-sperm cells (MP-SC). Roughly 200 sections were generated for each, from which 5 µm samples were captured and determined to yield 0.05 - 0.1 ng of RNA, which is consistent with comparable cell types from prior gametogenic cell LCM studies (Okada et al., 2013). Sections were pooled into two biological replicates per LCM sample type, with RNA extracted from each duplicate sample independently using the Arcturus^®^ PicoPure^®^ RNA Isolation Kit (Applied Biosystems^TM^). Total RNA recovered per sample was resuspended to a total volume of 10 µL and subjected to two or three rounds of amplification using the MessageAmp II RNA amplification kit (Ambion), as per the manufacturer’s instructions.

To enrich the collection of nuclei from generative and sperm cells of mature pollen, anthers and stigmas were collected from flowers at anthesis and subjected to osmotic shock in Brewbaker and Kwack medium (Brewbaker and Kwack, 1963), pH 6.5, supplemented with 12.5% (w/v) sucrose in 15 mL centrifuge tubes. The homogenate was mixed on a shaker at 130 rpm for 30 min at room temperature, filtered through 150 µm and then 30 µm CellTrics nylon sieves (Partec GmbH), and collected in 2 mL microfuge tubes. The homogenate was centrifuged at maximum speed for 2 min to pellet cells, and 50 µL was layered on 0.5 mL of 10% Percoll in 2 mL microfuge tubes that were centrifuged at 900 × g for 2 min. Next, 1.5 mL of 0.52 M (10%) mannitol and 10 mM MOPS buffer (pH 7.5) solution was added, and the tubes were centrifuged for 2 min at maximum speed. From the resulting pellet, 20 µL was added to 30 µL of Arcturus^®^ PicoPure^®^ RNA isolation buffer and snap frozen in liquid nitrogen before storing at -80°C. RNA was extracted from the generative and sperm nuclei samples and amplified as described for the LCM samples.

### Library preparation and RNA sequencing

Transcriptome libraries were prepared from amplified RNA using the Illumina Truseq mRNA protocol (Australian Genome Research Facility, Australia). RNA sequencing did not include poly-A purification or fragmentation of RNA, instead proceeding directly to adapter annealing and first-strand cDNA synthesis. Duplicate libraries were divided among four lanes of a HiSeq2500 flow cell and sequenced to generate paired-end sequence reads 125 bp in length for each end (2 × 125 bp reads).

### Cowpea genome annotation transfer

To compare gene expression between two cowpea varieties (IT97K-499-35 and IT86D-1010), reads mapping against their own genome were considered. IT86D-1010 whole-genome annotation was made consistent with the IT97K-499-35 gene ID system. For genome annotation transfer from IT97K-499-35 (PacBio sequencing, genome annotated by Phytozome) to IT86D-1010 (genome generated using Oxford Nanopore Technology (ONT) long read sequencing, downloaded from CSIRO: https://data.csiro.au/collection/csiro:63431), the repetitive and transposable elements, including interspersed repeats and low-complexity DNA, were identified by HiTE (Hu et al., 2024), with --thread 31, --plant 1, --recover 1, --domain 1, --annotate 1, --BM_RM2 1, --BM_EDTA 1, --BM_HiTE 1 parameters. This was followed by whole-genome soft masking by RepeatMasker (https://www.repeatmasker.org/), with -norna,-no_is, -xsmall, -gff, -lib confident_TE.cons.fa parameters. Next, annotation transfer between the two masked genomes using liftoff (Shumate et al., 2021) was performed with -p 31, -polish, -cds, -a 0.95, -s 0.9, -exclude_partial, -u unmapped_features.txt settings. The transfer rate was calculated by comparing annotation files between IT97K-499-35 and IT86D-1010 under Python.

### RNAseq read mapping and quantification

STAR (version 2.2.1; Dobin et al., 2013) ‘--runMode genomeGenerate’ was used to generate mapping indexes of the cowpea IT97K-499-35 genome (Lonardi et al., 2019) accessed from Phytozome 13 (https://phytozome-next.jgi.doe.gov/; Goodstein et al., 2012) and re-annotated IT86D-1010 genome with ‘--genomeSAindexNbases 13 --sjdbOverhang 100’ parameters. Paired-end raw reads from LCM and whole tissue datasets had their adapters removed and low-quality reads were filtered, with 15 random/low-quality nucleotides at the 5’-end trimmed using Fastp (Chen et al., 2018). Reads shorter than 40 bp were discarded using ‘-w 11 -l 40 -f 15 -F 15’ parameters. Remaining trimmed reads were mapped to the IT97K-499-35 or IT86D-1010 genome based on the sample genotype using STAR (Dobin et al., 2013) with ‘--twopassMode Basic --outFilterScoreMinOverLread 0.33 --outFilterMatchNminOverLread 0.33 --outSAMstrandField intronMotif --outSAMtype BAM SortedByCoordinate --quantMode GeneCounts --outReadsUnmapped None --limitBAMsortRAM 25000000000 -- limitOutSJcollapsed 1000000 --alignMatesGapMax 500 --alignIntronMax 20000’ settings. This allowed unpaired alignments and discard discordant aligned paired reads that are not compliant with the forward/reverse alignment of both mates to be removed. Following sorting by leftmost coordinates, BAM files generated were then converted to index files using the Samtools index (Li et al., 2009). The FeatureCounts function from Rsubread (Liao et al., 2019) was used to perform read count quantification, and transcripts per million (TPM) values were calculated using default settings. Read count and TPM files were merged, then gene length-corrected trimmed mean of M values (GeTMM) normalisation (Smid et al., 2018) was performed for cross-normalisation between and within samples.

### Visualisation of GeTMM normalisation and t-SNE sample clustering

To confirm TPM sample quality following GeTMM normalisation, RLE plots were generated through plotRLE() Function from EDASeq Package (Risso et al., 2011) in R. Each sample generated a boxplot of all deviations from its median expression. After normalisation, t-distributed stochastic neighbour embedding (t-SNE) projection and clustering analysis were performed to infer the sample relationship. To better demonstrate closely related samples and high-dimensional RNAseq data on a 2D plot, the Rtsne package (Van Der Maaten, 2014; Van der Maaten and Hinton, 2008) was used in R, with 100 dimensions in the initial PCA calculation (initial_dims = 100), 50000 iterations (max_iter = 50000), high accuracy (theta = 0.003), dims = 2, check_duplicates = TRUE, pca = TRUE, partial_pca = FALSE, verbose = getOption(“verbose”, FALSE), is_distance = FALSE, Y_init = NULL, pca_center = TRUE, pca_scale = FALSE, normalize = TRUE, momentum = 0.5, final_momentum = 0.8, eta = 200, exaggeration_factor = 12, and other parameters set as default, except perplexity = 2 for LCM samples and perplexity = 3 for whole tissue samples. Data visualisation was achieved using ggplot2 in R.

### Development of the VigExp visualisation platform

An interactive web-browser interface, called VigExp (*Vigna* Expression), was developed to visualise the transcriptome data and make it more user-friendly. The application was developed using open-source R packages, including shiny(https://github.com/rstudio/shiny), shinythemes (https://github.com/rstudio/shinythemes), DT (https://github.com/rstudio/DT), shinycssloaders (https://github.com/daattali/shinycssloaders), readr (https://github.com/tidyverse/readr), dplyr (https://github.com/tidyverse/dplyr), tibble (https://github.com/tidyverse/tibble), xml2 (https://cran.r-project.org/web/packages/xml2/index.html), rsvg (https://github.com/ropensci/rsvg), ComplexHeatmap (https://github.com/jokergoo/ComplexHeatmap), circlize (https://github.com/jokergoo/circlize), ggplot2 (https://github.com/tidyverse/ggplot2), ggpubr (https://github.com/kassambara/ggpubr), gridExtra (https://cran.r-project.org/web/packages/gridExtra/index.html), and cowplot (https://github.com/wilkelab/cowplot).

An electronic Fluorescent Pictograph (eFP) browser function for the cowpea transcriptomes was also developed using R. This enables the quick searching of genes of interest using a gene ID or keyword to visualise expression across different tissues and cell types. The platform integrates cowpea gene annotations from Phytozome 13 (https://phytozome-next.jgi.doe.gov/; Goodstein et al., 2012) and, based on OrthoFinder (Emms et al., 2019; Emms et al., 2026) homolog search against *Arabidopsis thaliana* (https://www.arabidopsis.org/), provides the top *Arabidopsis* gene ID hit based on E-value (E-value ≥ 50 very high quality, 10 ≤ E-value ≤ 50 high quality, E-value ≤ 10 moderate quality). It incorporates the full *Arabidopsis* gene name, primary gene short name, symbols, and gene function description from the TAIR database (https://www.arabidopsis.org/). In addition, transcription factor encoding genes were predicted using PlantTFDB (http://planttfdb.gao-lab.org/prediction.php; (Jin et al., 2017).

### RT-qPCR and identification of whole tissue, cell-type specific, and reference genes

A total of 26 tissue types, including whole root, root tip, mature root without root tip (Mature root), stem, young unexpanded leaf (YL), mature leaf (ML), pooled young flower bud (gynoecia <1.75 mm) (YFB Stage 1), pooled ovary (1.75 mm < gynoecia < 4 mm) (Ovary Stage 2), pooled ovary (5 mm < gynoecia < 8 mm) (Ovary Stage 3), ovule-mature embryo sac (O-MES), ovule-10 hour after pollination (O-10 hap), seed-3 dap (S-3 dap), seed-4.5 dap (S-4.5 dap), seed-6 dap (S-6 dap), seed-8 dap (S-8 dap), seed-10 dap (S-10 dap), seed-14 dap (S-14 dap), pod-3 dap (P-3 dap), pod-4.5 dap (P-4.5 dap), pod-6 dap (P-6 dap), pod-8 dap (P-8 dap), pod-10 dap (P-10 dap), pod-14 dap (P-14 dap), pooled anther (1.75 mm < gynoecia < 4 mm) (Anther Stage 2), pooled anther (5 mm < gynoecia < 8 mm) (Anther Stage 3), and anther-mature pollen (Anther MP) from IT86D-1010 plants, with 4 biological replicates for each tissue, except 3 for root tip and mature root (102 samples in total), were used for reverse transcription-quantitative PCR (RT-qPCR). The samples were harvested, snap-frozen in liquid nitrogen and stored at -80°C. For O-MES and O-10 hap samples, over 300 ovules were harvested for each biological replicate. RNA was isolated using a Maxwell RSC with the Maxwell RSC Plant RNA Kit (Promega, Madison, WI, USA) following the manufacturer’s instructions. Genomic DNA elimination for all RNA samples was performed using ezDNase™ Enzyme (Thermo Fisher Scientific, USA). cDNA was subsequently synthesised using 2 μg total RNA in 40 μL reactions with SuperScript IV reverse transcriptase (Invitrogen, Waltham, MA, USA). RT-qPCR was performed using a CFX384 real-time system (Bio-Rad, Hercules, CA, USA) with Bio-Rad CFX Maestro 1.1 using Luna® Universal qPCR Master Mix, three technical replicates for each sample, run at 95°C for 2 min, followed by 40 cycles for 95°C 15 s and 60°C 15 s. Melting curve was generated from 60°C to 95°C, increasing 0.5°C per cycle. Relative expression was normalised against the reference genes reported below, with gene-specific primers listed in Supplemental Table 2. Data visualisation and statistics were conducted using R or GraphPad Prism 11 with the 2 ^-ΔCt^ method.

To identify whole tissue and cell-type specific genes, expression was analysed using a log-transformed log2(TPM+1) matrix through an RNentropy (Zambelli et al., 2018) and TAU (Tissue specificity index, Yanai et al., 2005) calculation in R. Samples were screened using a TPM >2, Benjamini & Hochberg multiple test correction method, with a global p-value ≤0.05 and a local p-value ≤0.05 or TAU score ≥0.9. Results were visualised by heatmap generated through ComplexHeatmap (Gu et al., 2016) version 2.12.1 or the VigExp ‘Cowpea Easy Heatmap’ function.

To discover suitable reference genes for use in RT-qPCR studies, eight candidates were manually identified across the whole tissue transcriptome datasets based on expression level and standard variance using a log2(TPM+1) of CV (coefficient of variation) ≤ 0.1. Primers for each were subsequently designed (Supplemental Table 2) and tested against the whole tissue samples generated here for RT-qPCR (26 tissues, *n* = 3 - 4 biological replicates each). Findings were then assessed with RefFinder (Xie et al., 2023) plus R package ctrlGene using a calculation method of geNorm (Vandesompele et al., 2002) and NormFinder (Andersen et al., 2004) to rank the stability and reliability of each candidate.

### *In situ* hybridisation

*In situ* hybridisation (ISH) was performed as previously described (Jackson, 1991), with modifications for improved resolution of reproductive organs of cowpea. For sectioned specimens, mature unpollinated ovules were fixed in 4% paraformaldehyde and embedded in Paraplast. Sections of 12 μg thickness were attached to ProbeOnPlus slides (Fisher Biotech) and processed as previously described (Jackson, 1991). For whole-mount samples, developing ovule primordia, unpollinated mature ovules, and mature anthers were fixed in paraformaldehyde (4% paraformaldehyde, 2% Triton, and 1 × PBS in diethylpyrocarbonate DEPC-treated water) for 2 h at room temperature with gentle agitation, washed three times in 1 × PBS-DEPC water, and embedded in 15% acrylamide: bisacrylamide (29: 1) using precharged slides (Fisher Probe-On) treated with poly-l-Lys (Bass et al., 1997). Fixed pollen grains were stripped from anthers prior to embedding, and incubated in an enzymatic solution containing 1% driselase, 0.5% cellulose, and 1% pectolyase for 30 min in a humid chamber at 37°C. All samples were subsequently incubated in a 0.2 M HCl solution for 30 min at room temperature and in 1 µg/µL proteinase K for 30 min at 37°C, before processing as previously described (Garcia-Aguilar et al., 2005). Slides containing paraffin sections were mounted on Cytoseal, whereas slides containing whole specimens were mounted in either 50% (ovules) or 20% (pollen grains) glycerol, and analysed using a Leica DMRB microscope under Nomarski illumination.

### Generation of binary constructs, stable cowpea transformation, and CRISPR gene editing

To test the activity of cowpea promoters identified in this study, transformation vectors were developed using Golden Gate cloning. A 2,584 bp *VuEF1a* promoter and 730 bp *VuEF1a* terminator were amplified from the cowpea variety IT86D-1010 and combined with a gene encoding spectinomycin resistance for selecting positive transformation events, generating a final cassette of *pVuEF1a*:: *CTP-SpcN*:: *tVuEF1a* or *pVuEF1a*:: *CTP-aadA1*:: *tVuEF1a* for transgenic cowpea selection. To establish *VuUBQ4* and *VuUBQ10* promoter activity, a 1,599 bp *VuUBQ4* promoter, 2,610 bp *VuUBQ10* promoter, and 466 bp *VuHSP17.6* terminator were cloned from IT86D-1010 by PCR to generate *pVuUBQ4*:: *ZsGreen*:: *tVuHSP17.6* and *pVuUBQ10*:: *ZsGreen*:: *tVuHSP17.6* cassettes, respectively. To identify the *TAM* homolog in cowpea, *VuTAM2*, the protein domain of AtTAM and SlTAM predicted from NCBI containing a Cyclin N-terminal and Cyclin C-terminal domain were selected. HMMER was used with Pfam Cyclin_N and Cyclin_C hidden Markov models against all IT97K-499-35 protein sequences, with 43 proteins containing both domains being captured. Protein sequence alignment was performed using MAFFT (E-INS-i algorithm, scoring matrix 200 PAM / k = 2, and gap open penalty 1.53), including *Arabidopsis*, tomato, and rice TAM family sequences, and a phylogenetic tree was generated in Geneious using FastTree (Rate categories of sites 3,000). To perform a full gene knockout, two small guide RNAs (gRNA) TAM2_UQsg1 and TAM2_UQsg2 were designed by WU-CRISPR (Wong et al., 2015) and Geneious (off-target checking) to target the coding sequence of *VuTAM2* gDNA region close to the ATG start codon. The gRNA sequences were synthesised by Integrated DNA Technologies (IDT) and cloned into cowpea CRISPR vectors pUQHyG020b7 (*aadA1* selection marker) or pUQHyG020b1 (*SpcN* selection marker) by Golden Gate cloning (NEB, NEBridge Golden Gate Assembly Kit (*Bsa*I-HF v2)). All the key primers used for vector construction are listed in Supplemental Table 2. Cowpea transformation was performed according to Che et al. (2020) using the cowpea variety IT86D-1010. Gene editing of T1 and T2 plants was confirmed by Sanger sequencing of the desired cleavage site and analysed using ICE Analysis (https://ice.editco.bio/#/).

### Cytology, fluorescence microscopy and confocal microscopy

Transgenic shoots, stems, roots, flowers, shoot apical meristems, embryonic axes, cotyledons, and seeds were observed under a dissection fluorescence microscope with an Ex 480/30 excitation filter and 535/40 barrier filter. Exposure times were adjusted as appropriate for each sample, and IT86D-1010 wild-type tissue was used as a negative control to adjust the microscope settings and avoid detection of autofluorescence and overexposure. NIS-Elements software was used to capture images (Nikon). Leaf, pollen, and ovule samples were examined using a Zeiss LSM700 confocal microscope. Digital images were captured using ZEN black software (Carl Zeiss). ZsGreen was observed using a Zeiss 488 filter (20% power, Gain 670), with IT86D-1010 wild-type used as negative control. Propidium iodide-stained images were observed under Zeiss 555 filter (4% power, Gain 574). Images were processed using ZEN5.0 software and assembled in Adobe Illustrator. Dyads or tetrads were stained with 0.05% toluidine blue solution. Pollen viability was conducted using Alexander’s staining. Ovules stained with propidium iodide and clearing methods were processed according to Salinas-Gamboa et al. (2016) for samples requiring enhanced cell visualisation. The ClearSee (Kurihara et al., 2015) clearing method was used for visualising ZsGreen fluorescence signals in cowpea ovules.

### Development of a novel method for ploidy analysis, including ploidy verification of *VuTAM2* CRISPR knockout lines

Suitable reference genomes for determining cowpea and sorghum ploidy using flow cytometry were tested, including common bean (*P. vulgaris*), soybean (*G. max* var. Bunya), popping maize (*Z. mays* var. Everta), and sunflower (*H. annuus*). Nine paired combinations were tested in total. Seeds of the different species, as well as the *VuTAM2* knockout lines, were sown in 30-well trays, with young leaves from one- to two-week-old plants used for flow cytometry. CyStain UV Precise P Kit (containing Extraction Buffer and Staining Buffer) was used for sample preparation. Approximately 0.5 - 1 cm^2^ leaf tissue from cowpea/sorghum was mixed with a single reference species on a plastic Petri dish and 500 μL Extraction Buffer was added. The plant material was chopped finely with a razor blade for ∼1 min and incubated at room temperature for 1 min. The sample was then filtered through a 30 μm mesh filter (Sysmex) into a 3 mL glass tube (Sysmex) and 1 - 2 mL Staining Buffer was then added and allowed to incubate for 1 min. All samples were run on the CyFlow Space flow cytometer using FloMax 2.11 software with the Gain-value set to 551 for peak detection. The G_0_/G_1_ peaks Mean-x value for cowpea and soybean, or sorghum and maize were collected to calculate the ploidy level. Relevant formulas for cowpea and sorghum normalisation (Norm. factor, Supplemental Table 4 - 5) and ploidy calculation are as follows:

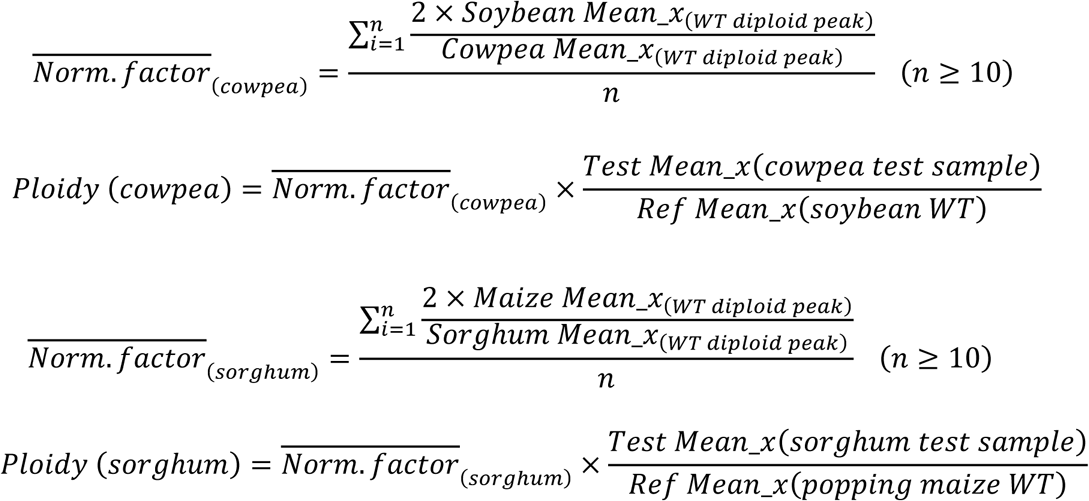

## ACKNOWLEDGMENTS

We are grateful to Tracy How and Dilrukshi Nagahatenna from CSRIO for their assistance with growing cowpea plants for LCM. Thanks also to Corteva Agriscience for providing the *CTP-SpcN* sequence, Peggy Ozias-Akins and Joann A. Conner from the University of Georgia for providing the *CTP-aadA1* and *ZsGreen* sequences, Nicolas Lester and Jacqui Foster of UQ for their assistance with stable transformations and pollen viability, and Guoquan Liu of UQ for providing haploid, diploid, and tetraploid Tx430 sorghum seeds for the method developed in this paper to verify ploidy.

## AUTHOR CONTRIBUTIONS

Conceptualization, H.S., A.M.G.K., and B.J.F.; methodology, H.S., D.M., N.G., M.R., M.J., S.D.J., J.H.Y., J.D., Y.L., A.M., G.L.-M, R.E.-G., R.S.-G., and I.A.-M.; formal analysis and data curation, H.S., A.M.G.K., and B.J.F.; supervision and project administration, J.-P.V.-C., A.M.G.K., B.J.F.; funding acquisition, A.M.G.K.; writing – original draft, H.S., A.M.G.K., and B.J.F.; writing – review & editing, H.S., J.-P.V.-C., A.M.G.K., and B.J.F.

## SUPPLEMENTAL INFORMATION

Correspondence and requests for materials should be addressed to Brett J. Ferguson.

---

**Supplemental Figure 1.**
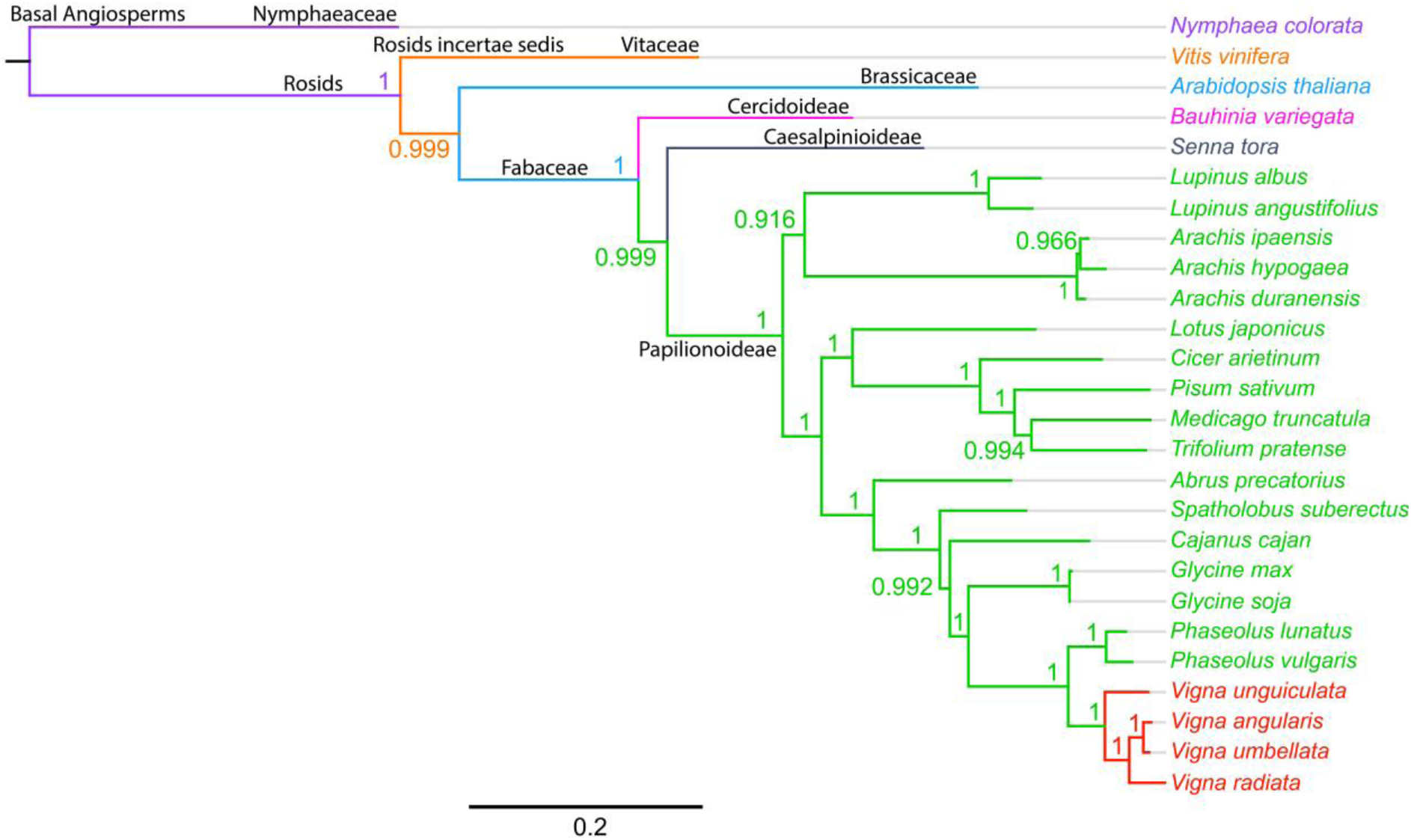
Phylogenomic tree of several legume species based on proteome sequence maximum-likelihood analysis.

**Supplemental Figure 2.**
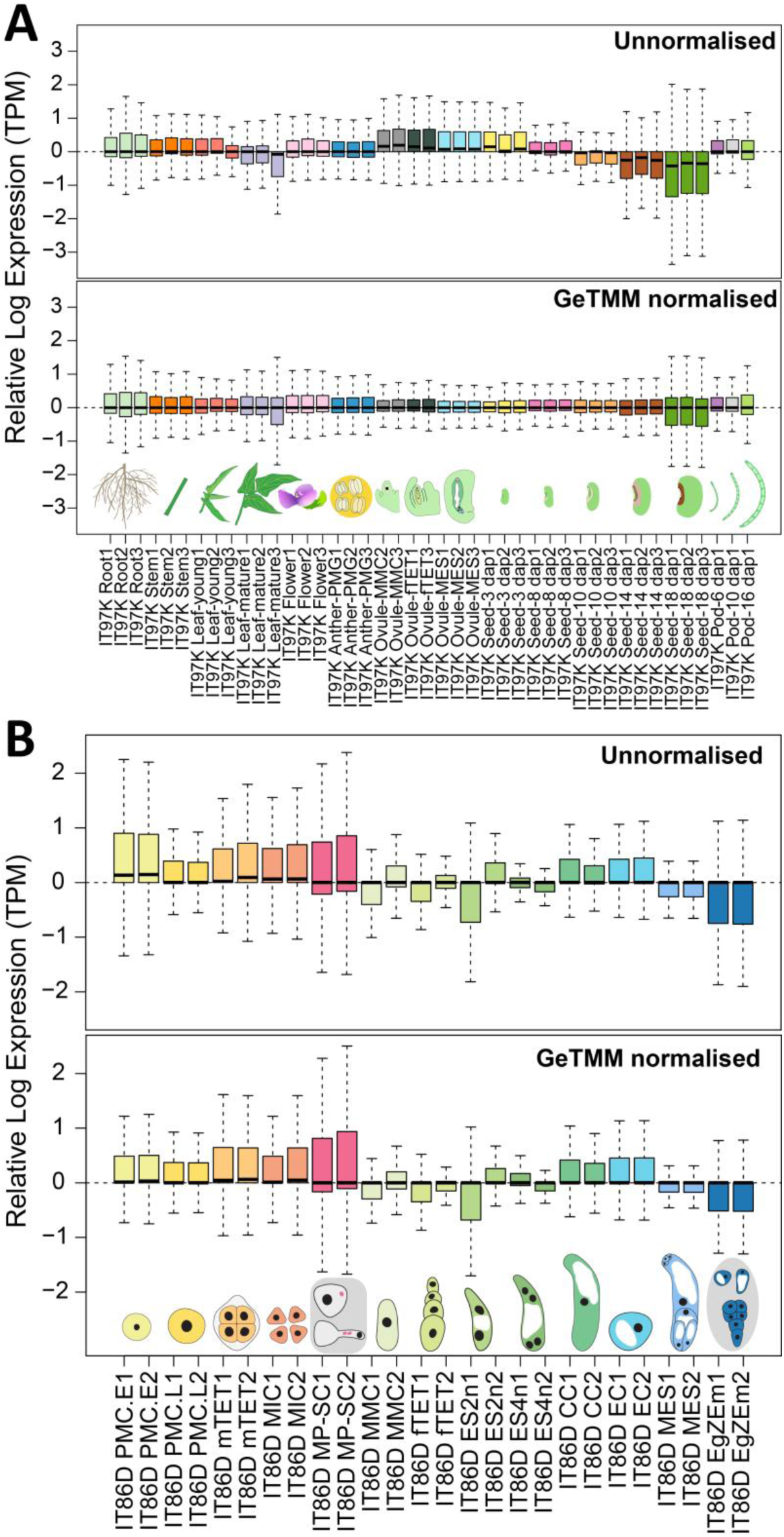
GeTMM normalisation for whole tissue and LCM RNAseq data sets.

**Supplemental Figure 3.**
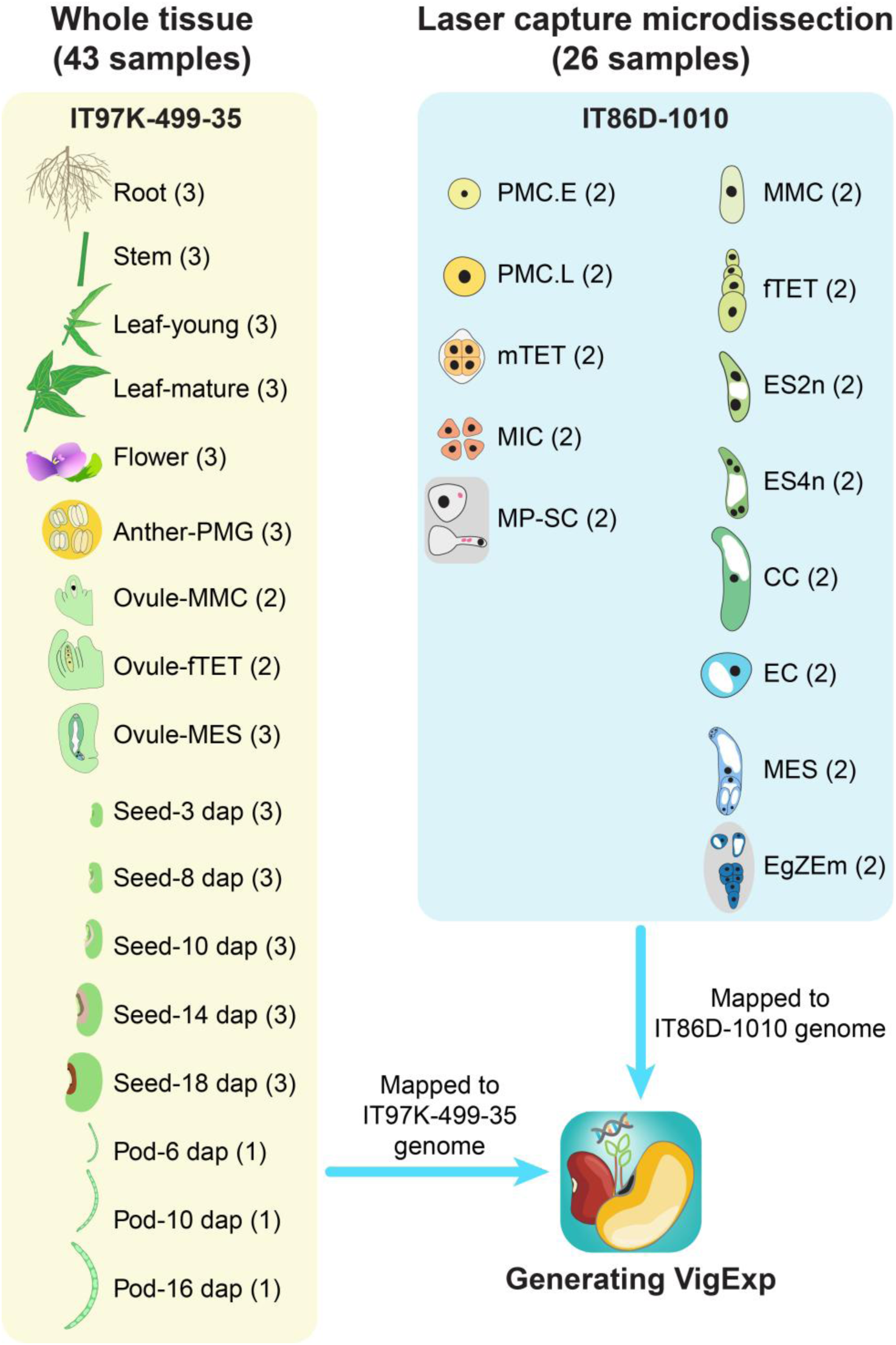
Overview of samples sourced for transcriptomic datasets reported in this study.

**Supplemental Figure 4.**
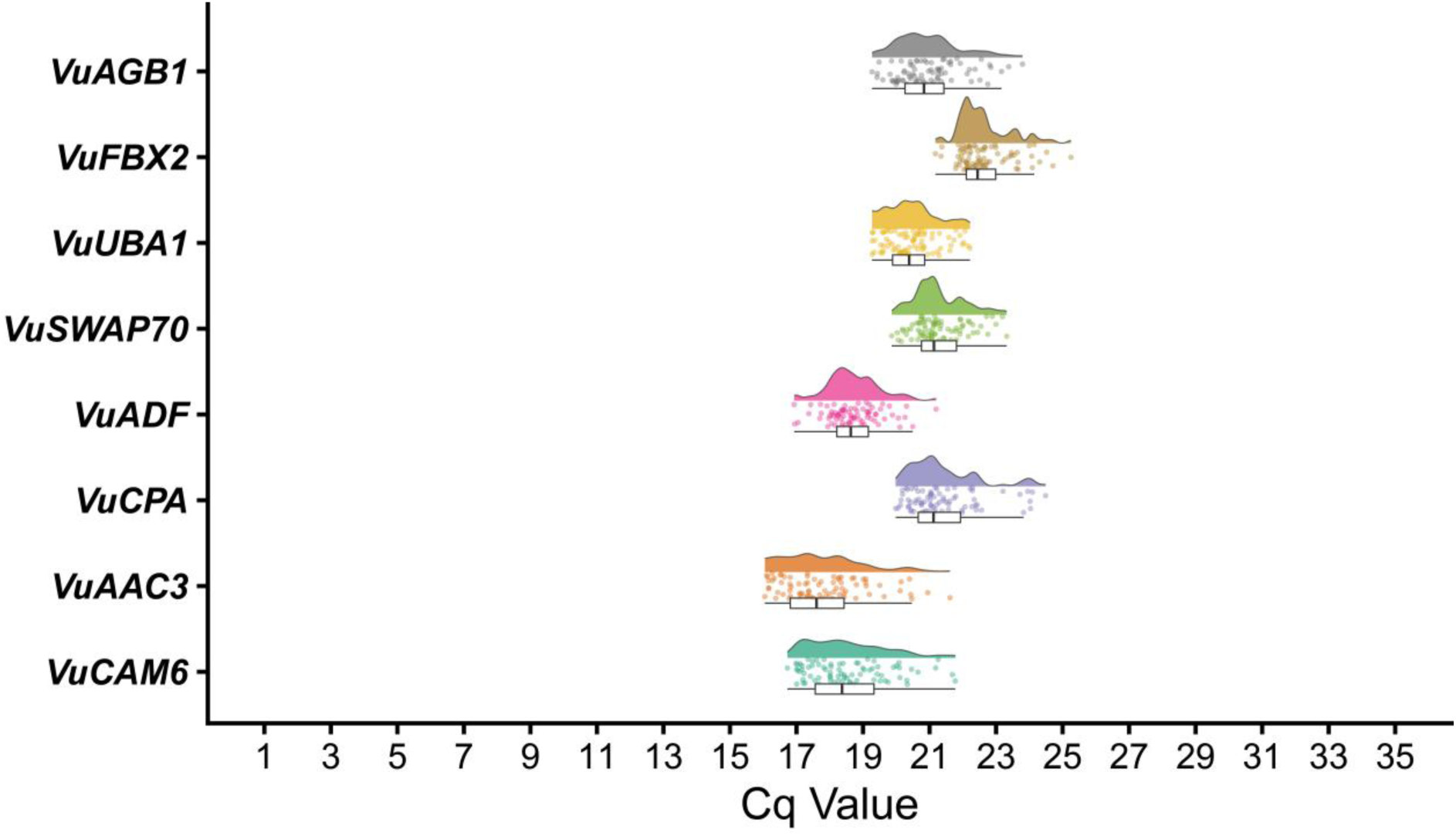
Cq stability of cowpea reference genes across 78 samples, including 26 sample types.

**Supplemental Figure 5.**
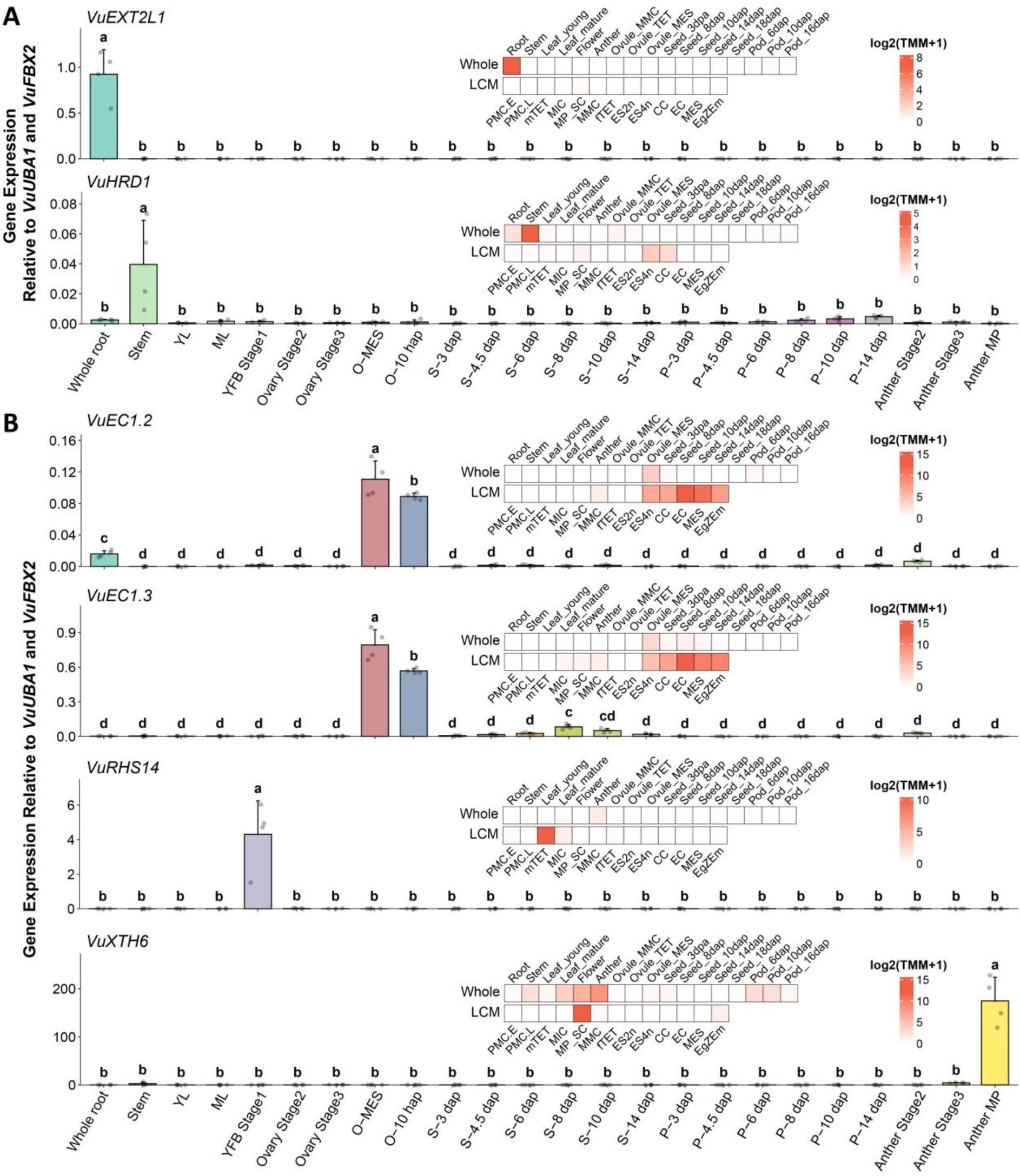
Whole-tissue and cell-type specific gene expression patterns determined via RT-qPCR to validate the transcriptomic datasets. **(A)** Whole-tissue specific genes identified using VigExp and then validated by RT-qPCR. **(B)**Cell-type-specific genes identified using VigExp and then validated by RT-qPCR.

**Supplemental Figure 6.**
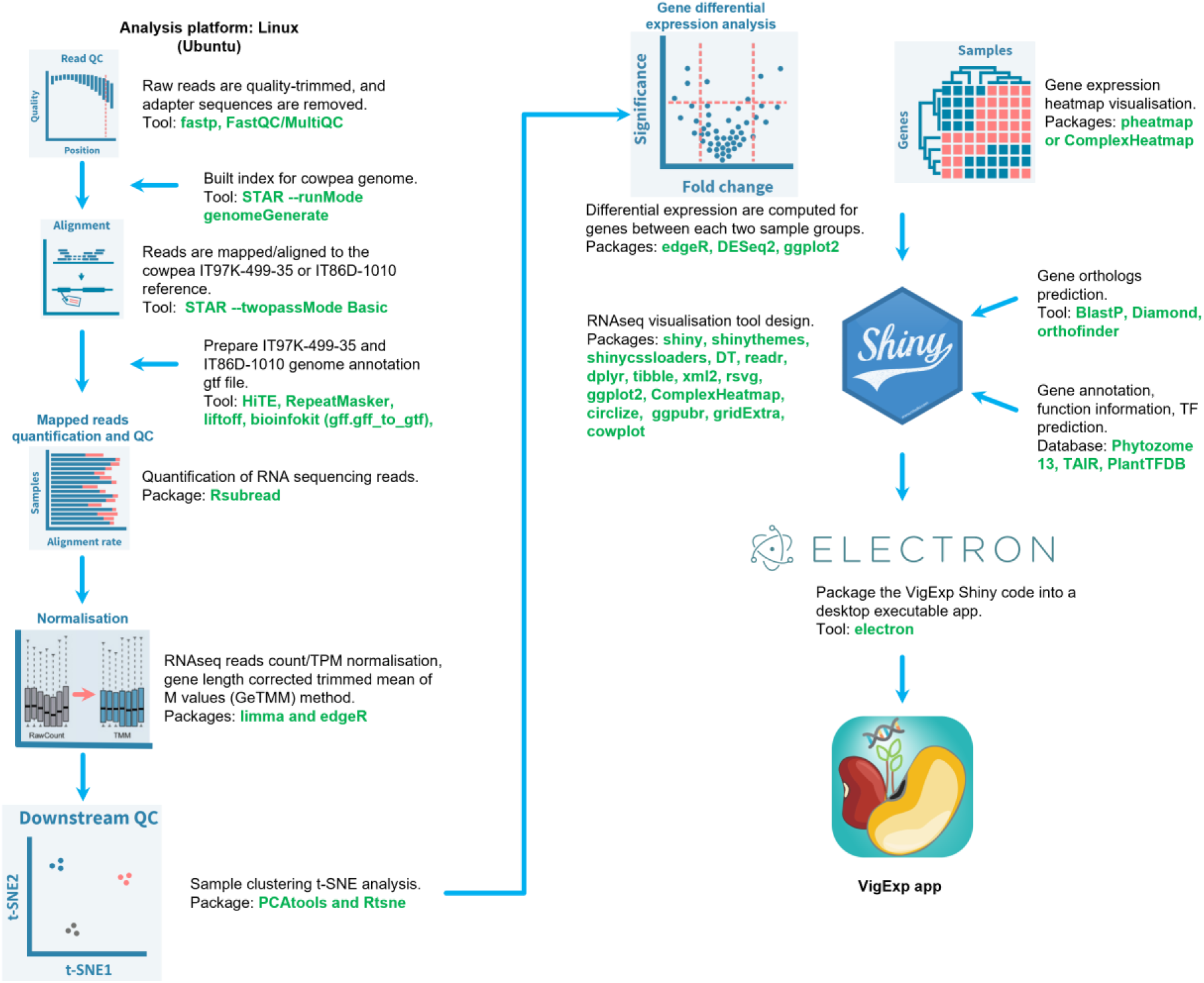
Flow chart for developing the VigExp platform.

**Supplemental Figure 7.**
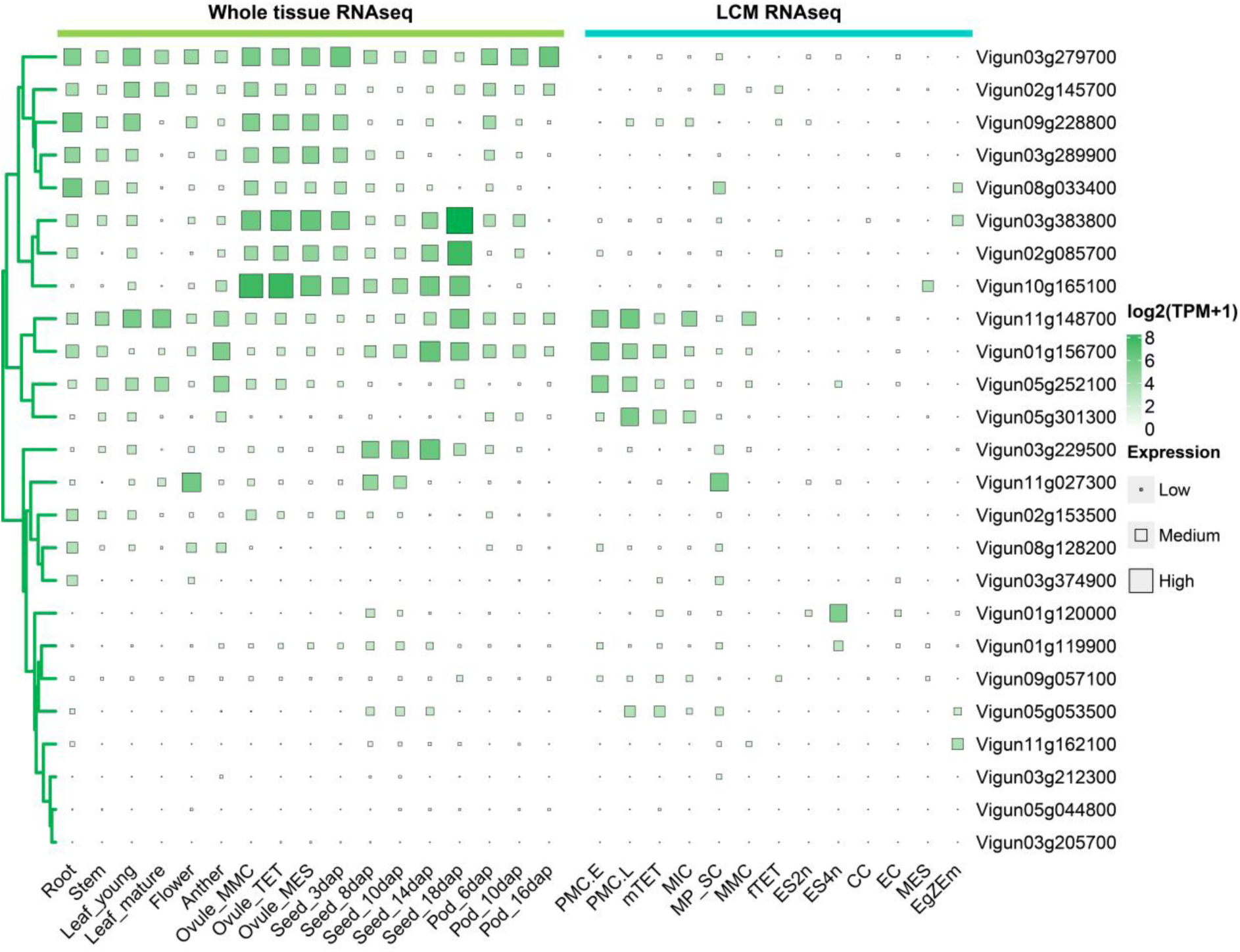
Heatmap of the cowpea *VuBBML* gene family (25 double AP2 domain proteins).

**Supplemental Figure 8.**
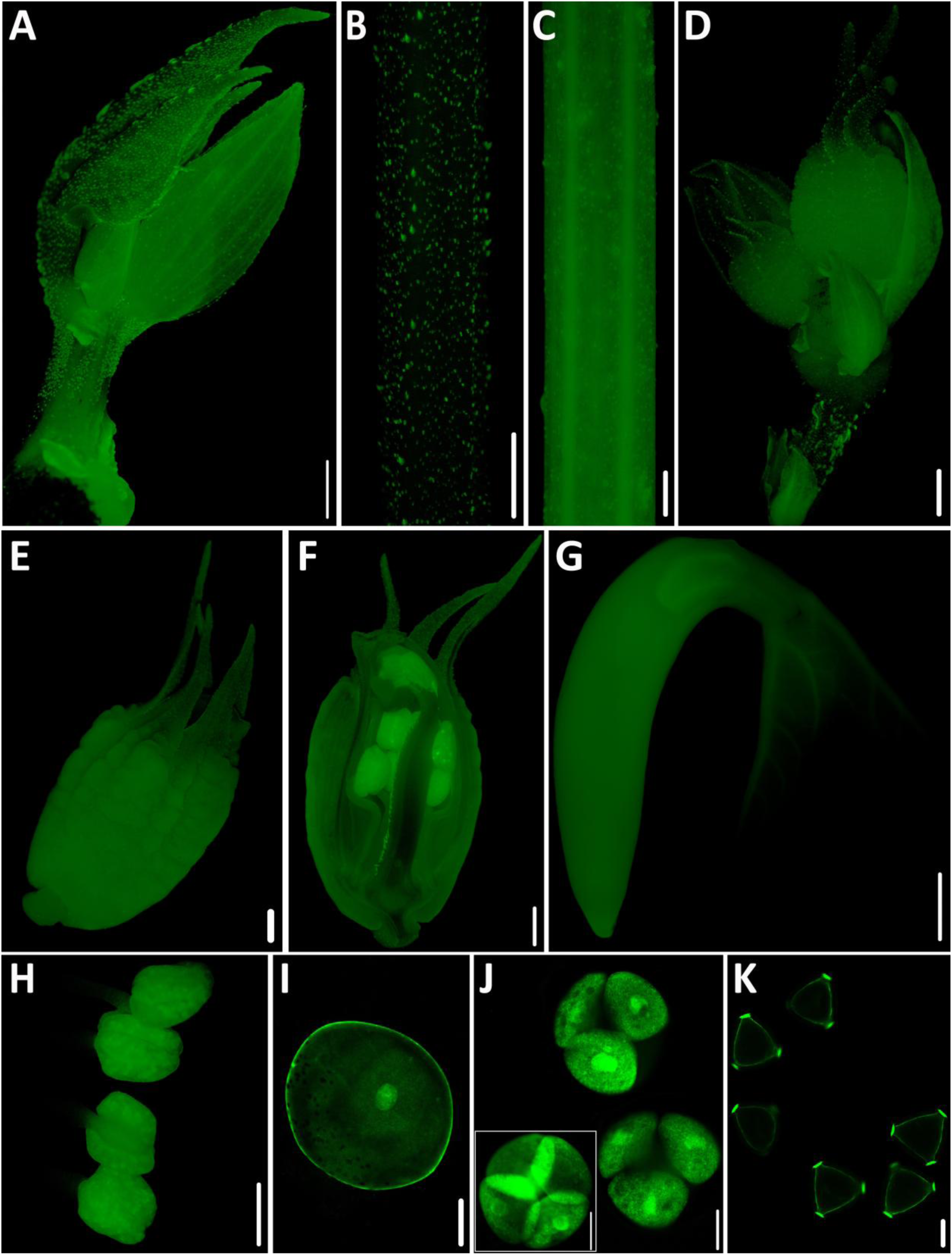
Cowpea *VuUBQ4* constitutive promoter activity in different cowpea tissues, including reproductive cells and stages. **(A)** *pVuUBQ4*:: *ZsGreen*:: *tVuUBQ4* expression in the young shoot, **(B)** young stem, **(C)** mature stem, **(D)** mature flower buds, **(E)** undissected and **(F)** dissected floral bud at stage 1F-VII, **(G)** embryonic axis, **(H)** fully-mature anther, **(I)** pollen mother cell, **(J)** tetrads, **(K)** microspores. Scale Bars: **A - H** = 1000 μm; **I - K** = 20 μm.

**Supplemental Figure 9.**
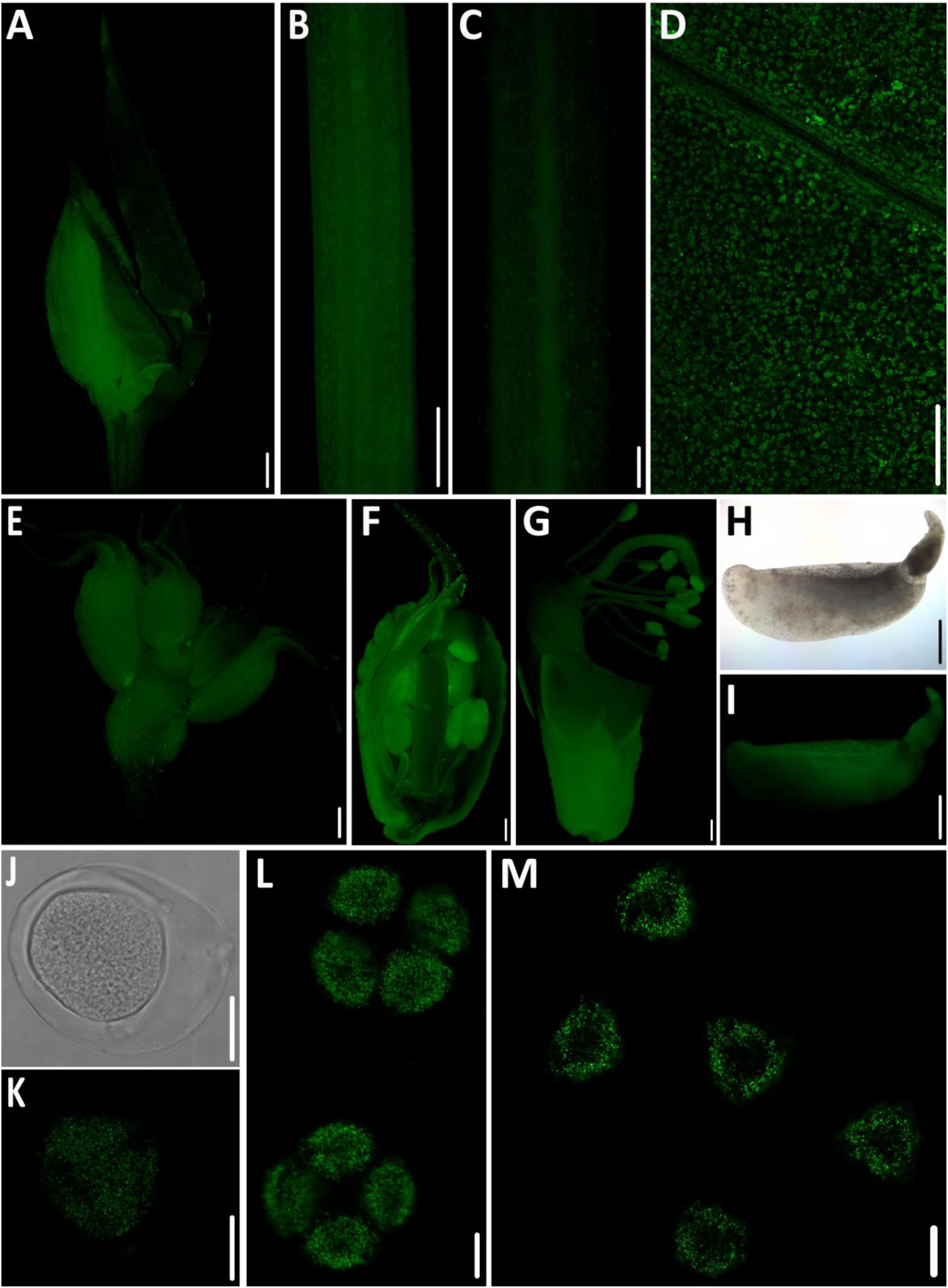
Cowpea *VuUBQ10* constitutive promoter activity in different cowpea tissues, including reproductive cells and stages. **(A)** *pVuUBQ10*:: *ZsGreen*:: *tVuUBQ10* expression in the young shoot, **(B)** young stem, **(C)** mature stem, **(D)** mature leaf, **(E)** mature flower buds, **(F)** dissected floral bud at stage 1F-VII, **(G)** dissected floral bud at stage 1F-IX, **(H)** dissected embryo at young cotyledon stage under bright field and **(I)** dark field, **(J)** pollen mother cell under bright field and **(K)** dark field, **(L)** tetrads, and **(M)** microspores. Scale Bars: **A - I** = 1000 μm; **J - M** = 20 μm.

**Supplemental Figure 10.**
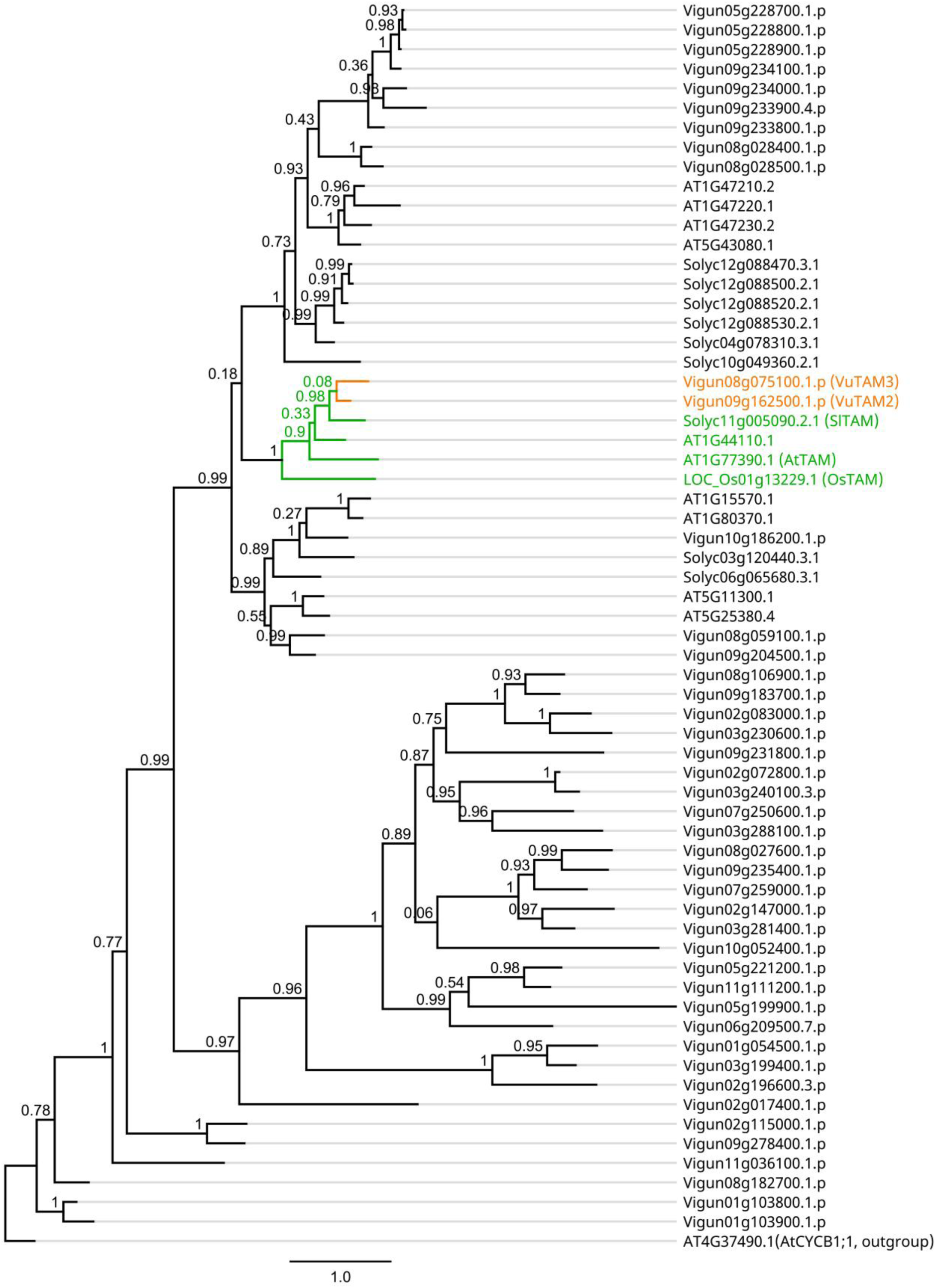
Cowpea protein phylogenetic tree to identify *VuTAM*.

**Supplemental Figure 11.**
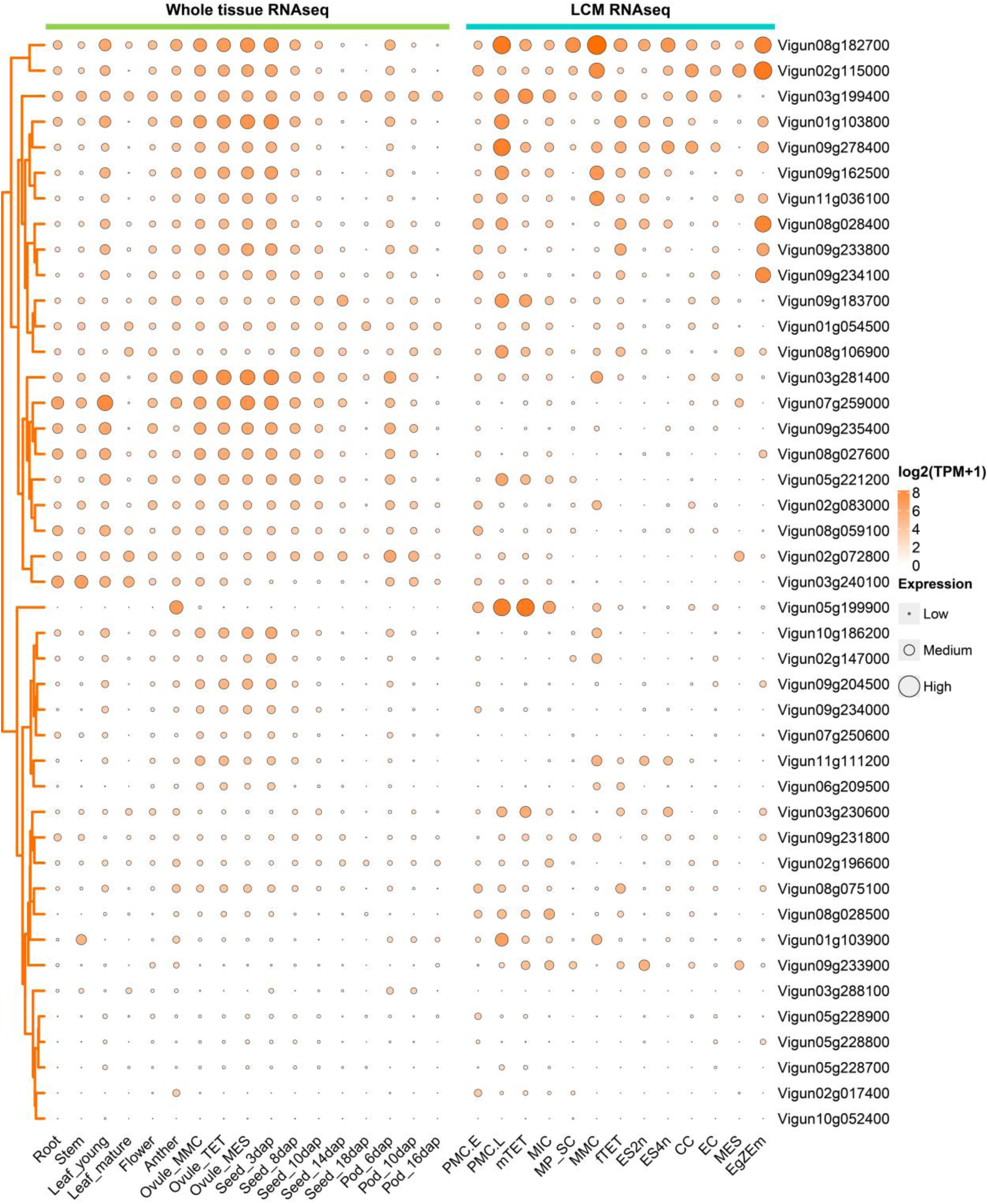
Heatmap of the cowpea *VuTAM* gene family (43 Cyclin N-terminal and Cyclin C-terminal domain proteins).

**Supplemental Figure 12.**
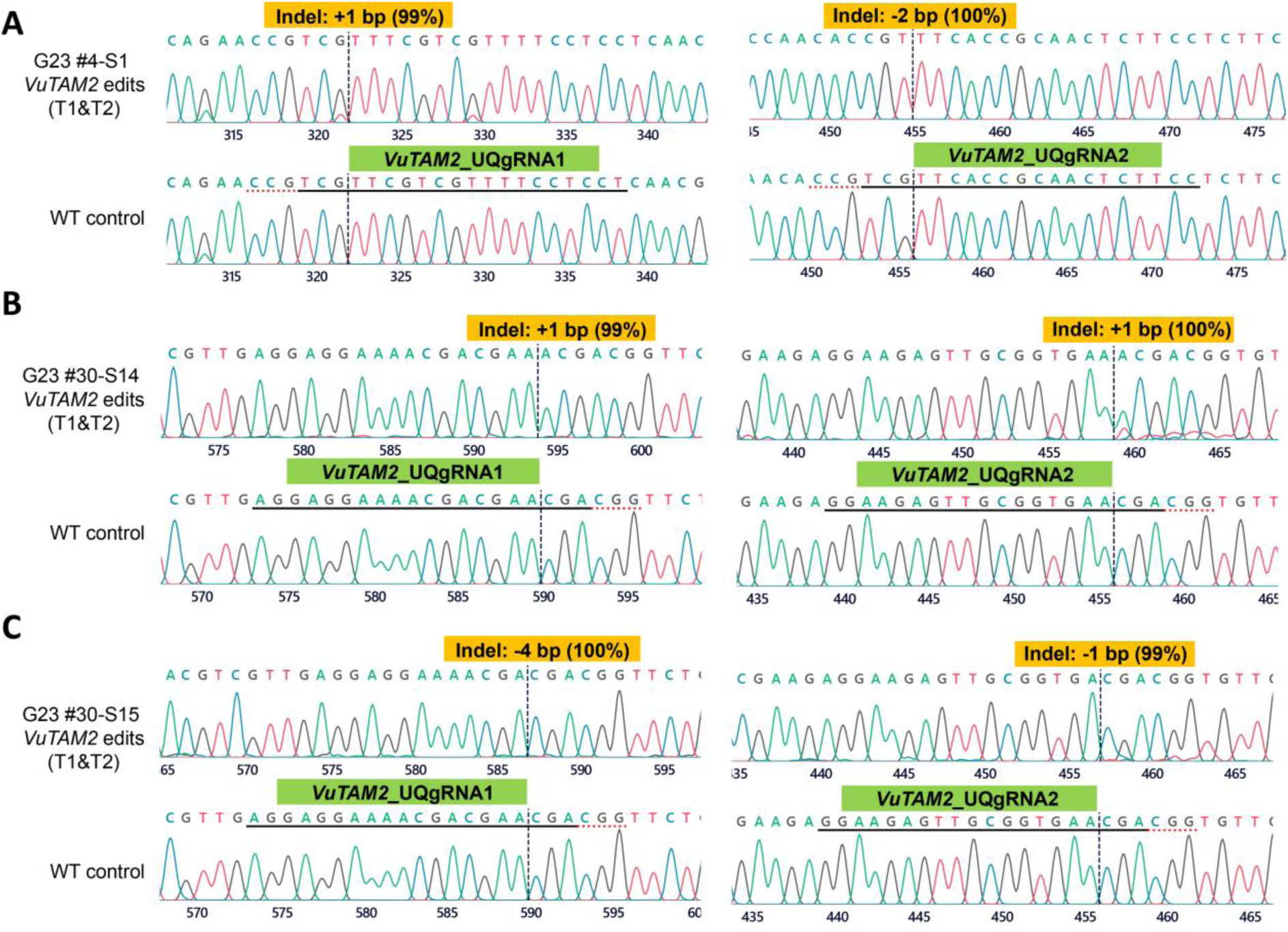
V*u*TAM2 CRISPR gene-edited biallelic knockout line sequencing results. **(A)** Line G23 #4-S1 has stable biallelic edits in the *VuTAM2* gene across T1 and T2 generations, with Sanger sequencing results exhibiting +1 bp at the UQgRNA1 and -2 bp at the UQgRNA2 cleavage sites. **(B)** Line G23 #30-S14 has stable biallelic edits in the *VuTAM2* gene across T1 and T2 generations, with Sanger sequencing results exhibiting +1 bp at the UQgRNA1 and +1 bp at the UQgRNA2 cleavage sites. **(C)** Line G23 #30-S15 has stable biallelic edits in the *VuTAM2* gene across T1 and T2 generations, with Sanger sequencing results exhibiting -4 bp at the UQgRNA1 and -1 bp at the UQgRNA2 cleavage sites. The horizontal black underlined region represents the guide sequence. The horizontal red underline is the PAM site. The vertical black dotted line represents the actual cut site.

**Supplemental Figure 13.**
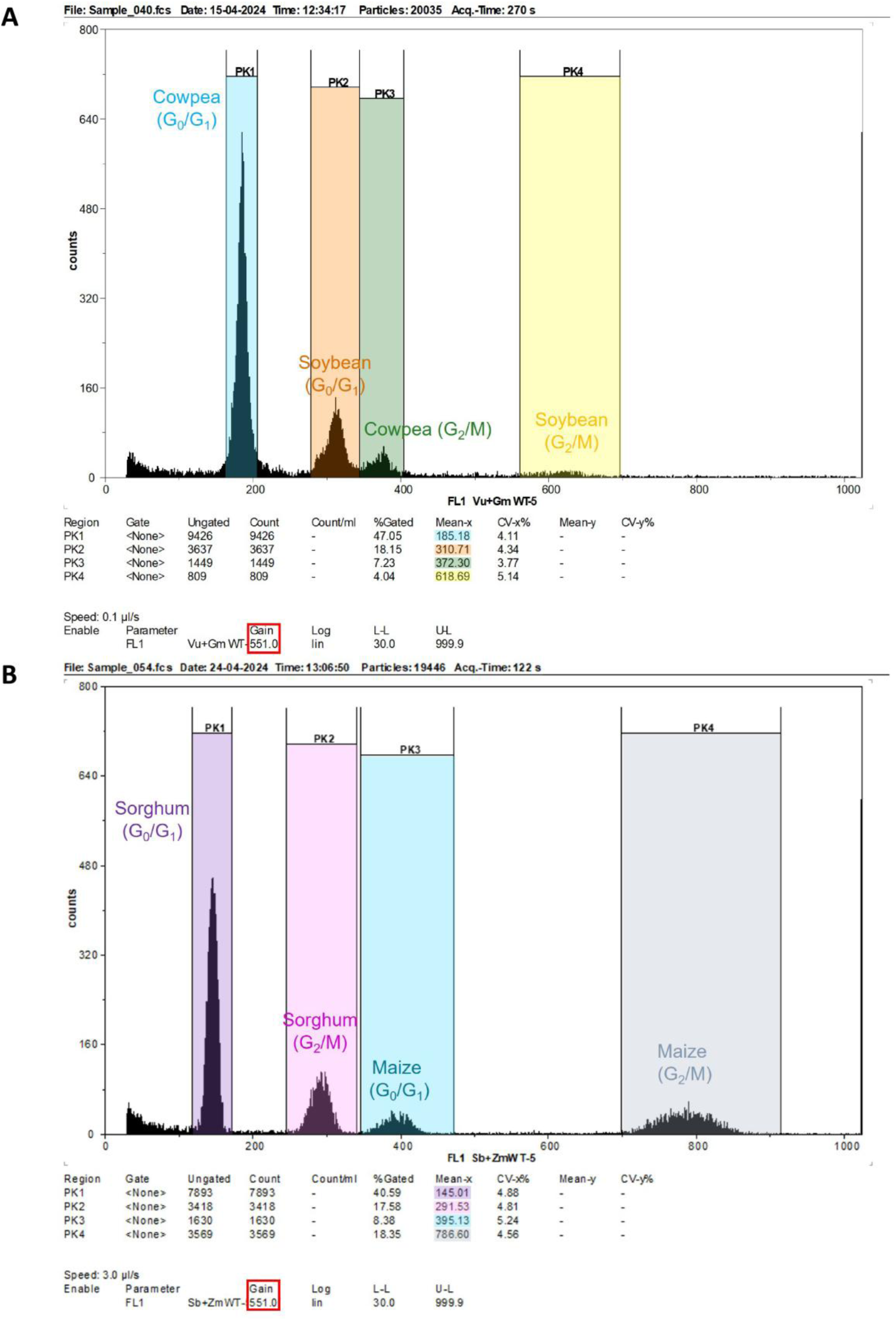
Cowpea/sorghum flow cytometry ploidy normalisation by using soybean/popping maize as a reference. **(A)** Cowpea ploidy level verification using wild-type soybean (*Glycine max* var. Bunya) as a reference. **(B)** Sorghum ploidy level verification using wild-type popping maize (*Zea mays* var. Everta) as a reference.

**Supplemental Figure 14.**
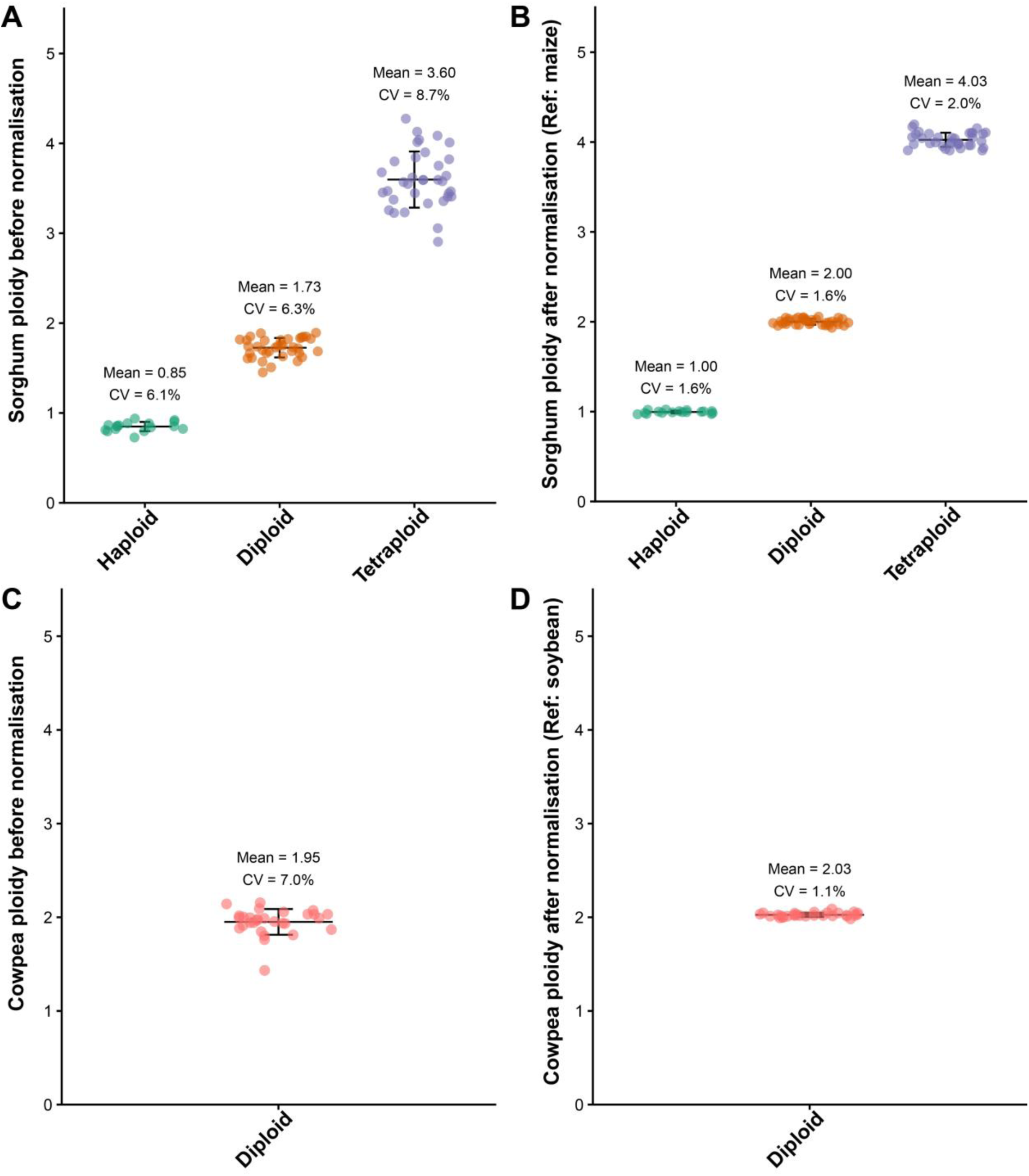
Accurate ploidy identification method tested in sorghum and cowpea. Haploid, diploid, and tetraploid Tx430 sorghum ploidy verification before **(A)** and after **(B)** normalisation. Diploid IT86D-1010 cowpea ploidy verification before **(C)** and after **(D)** normalisation.

Supplemental Table 1. Read mapping info summary for each reproductive cell-type and whole tissue transcriptome.

Note: this will be uploaded as a separate Excel file, called ‘Supp Table1. RNAseq reads mapping info summary.xlsx’

**Supplemental Table 2.**
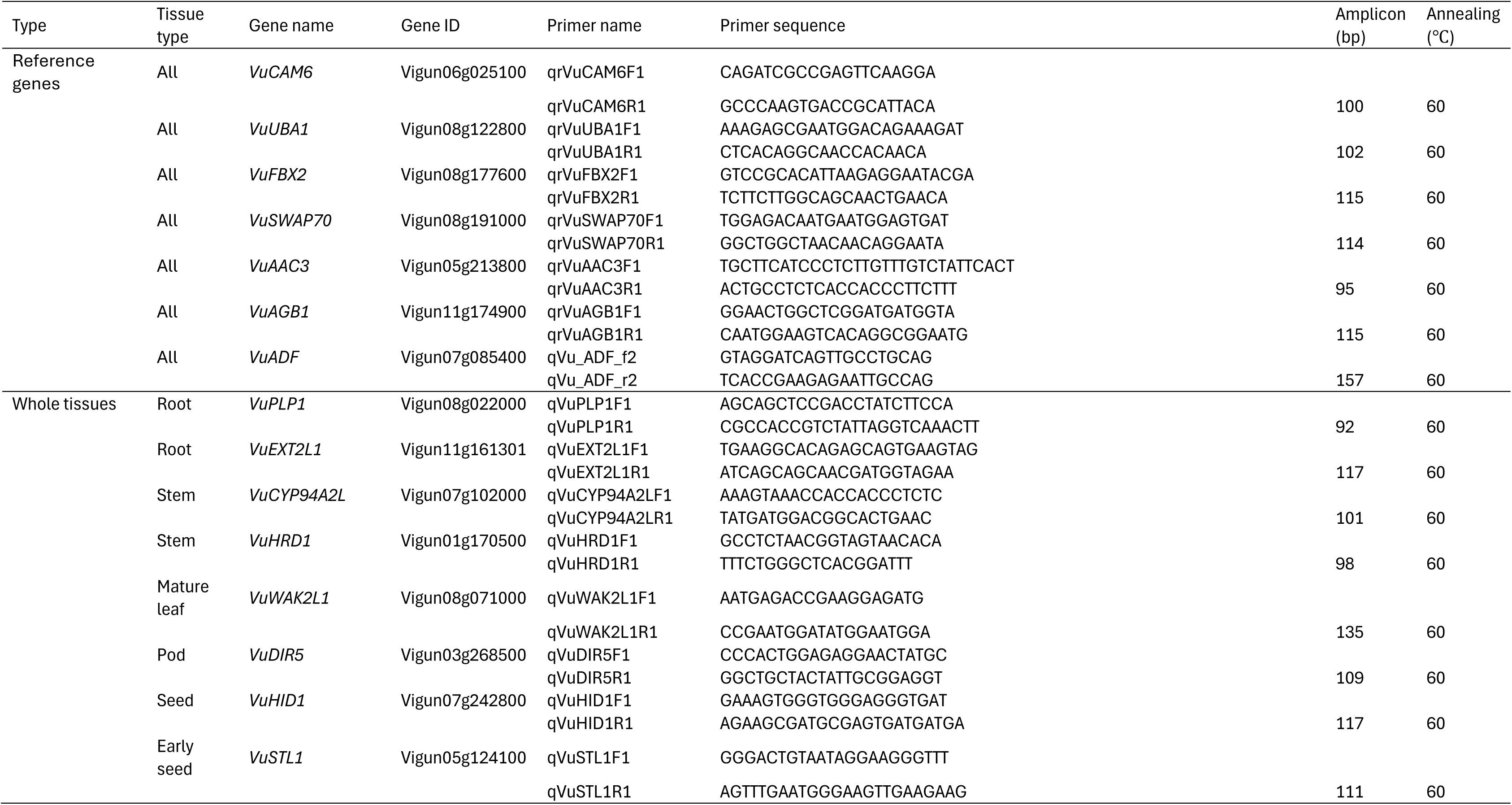
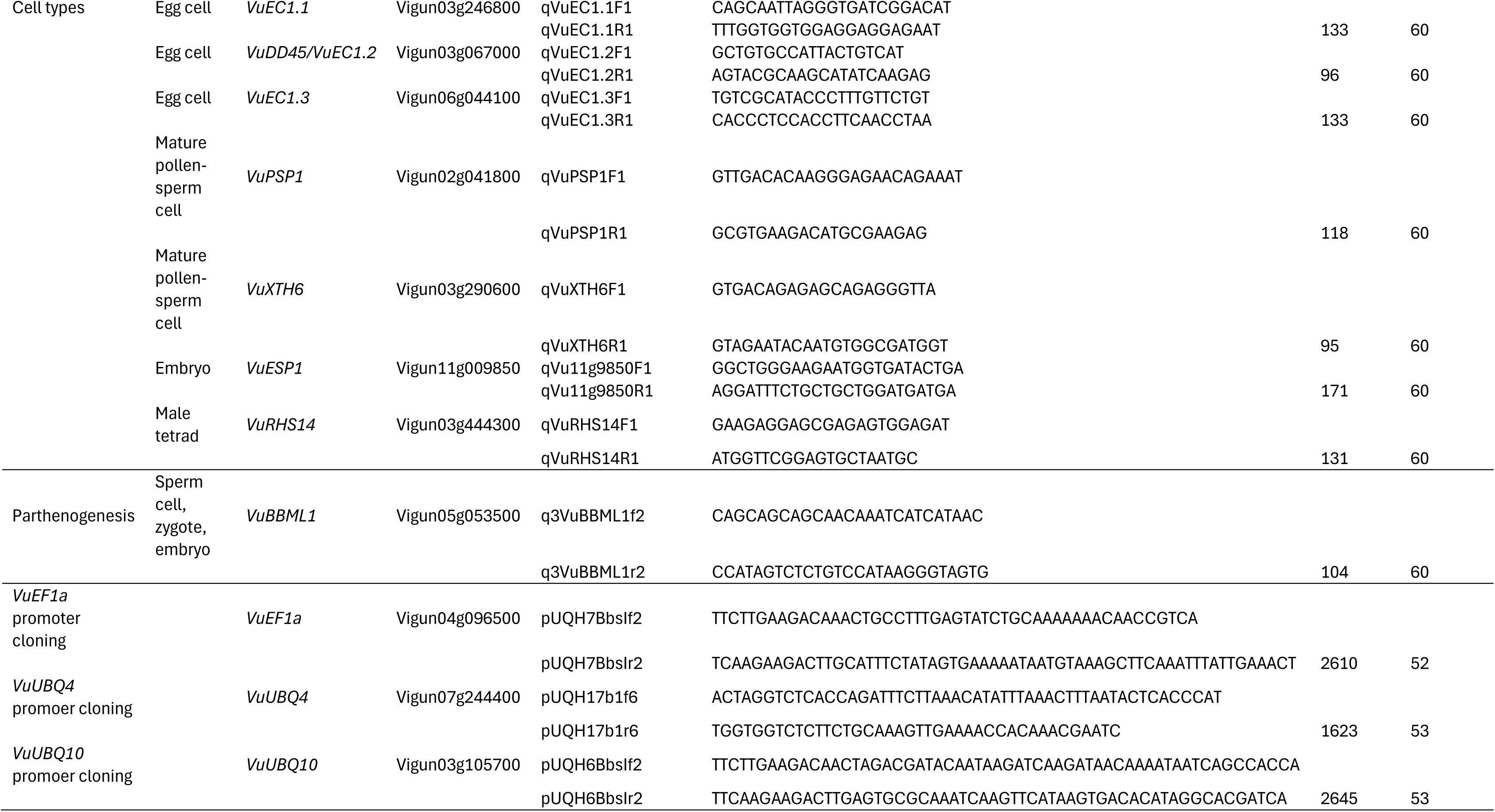
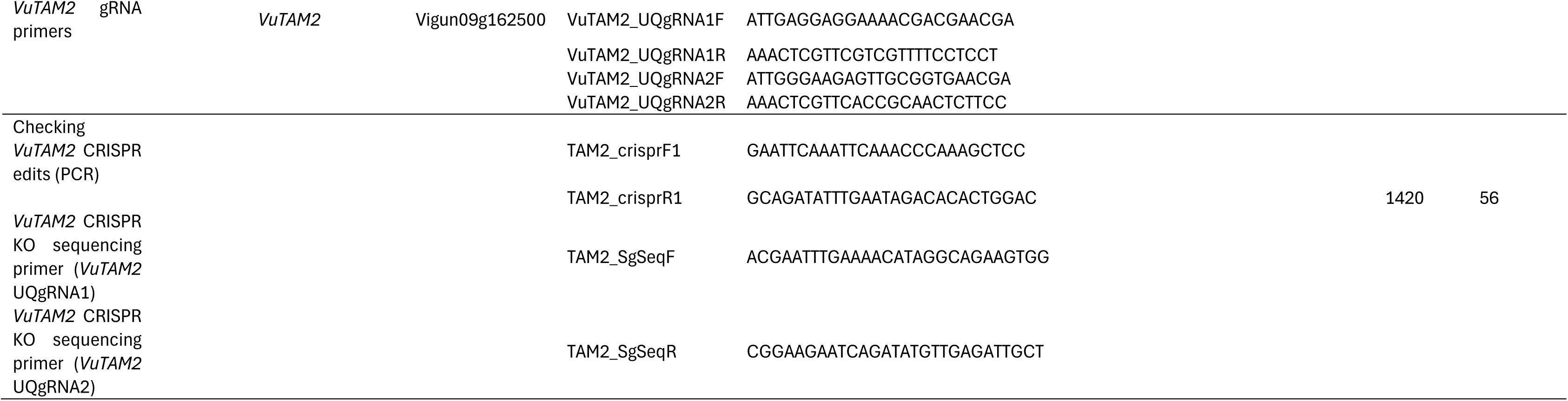
Key primers used in this study.

**Supplemental Table 3.**
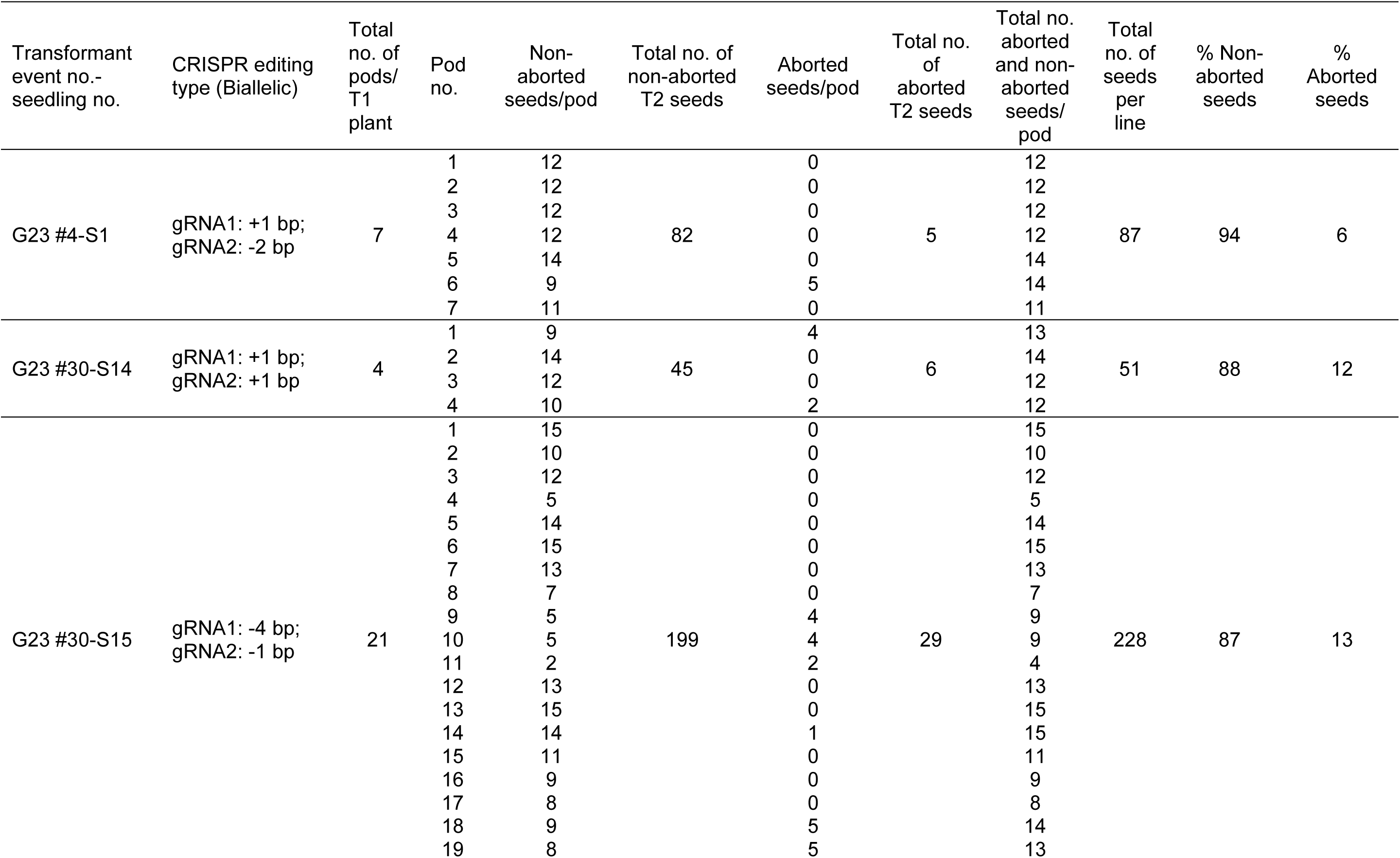
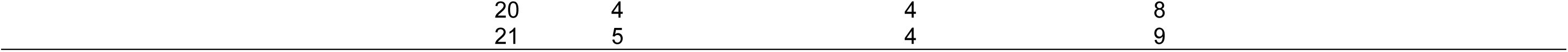
Pod and seed setting from *VuTAM2* knockout lines.

**Supplemental Table 4.** Cowpea ploidy normalisation factor calculation using wild-type soybean (*G. max* var. Bunya) as a reference.

| Sample ID | Gain | Cowpea Mean-x<br>(cowpea diploid peak<br>PK1) | Soybean Mean-x<br>(soybean diploid peak PK2) | Norm. factor |
| --- | --- | --- | --- | --- |
| Vu+Gm WT-1 | 551 | 168.46 | 285.62 | 3.39 |
| Vu+Gm WT-2 | 551 | 198.65 | 328.63 | 3.31 |
| Vu+Gm WT-3 | 551 | 180.26 | 303.21 | 3.36 |
| Vu+Gm WT-4 | 551 | 187.73 | 311.52 | 3.32 |
| Vu+Gm WT-5 | 551 | 185.18 | 310.71 | 3.36 |
| Vu+Gm WT-6 | 551 | 201.71 | 337.52 | 3.35 |
| Vu+Gm WT-7 | 551 | 199.32 | 328.48 | 3.30 |
| Vu+Gm WT-8 | 551 | 180.71 | 301.2 | 3.33 |
| Vu+Gm WT-9 | 551 | 199.18 | 332.96 | 3.34 |
| Vu+Gm WT-10 | 551 | 187.84 | 313.15 | 3.33 |
| Vu+Gm WT-11 | 551 | 199.15 | 330.29 | 3.32 |
| Mean value |  |  | *Norm. factor*<sub>(cowpea)</sub> | **3.34** |

**Supplemental Table 5.** Sorghum ploidy normalisation factor calculation using wild-type popping maize (*Z. mays* var. Everta) as a reference.

| Sample ID | Gain | Sorghum Mean-x<br>(sorghum diploid peak PK1) | Maize Mean-x<br>(maize diploid peak PK3) | Norm.<br>factor |
| --- | --- | --- | --- | --- |
| Sb+Zm WT-1 | 551 | 132.61 | 362.48 | 5.47 |
| Sb+Zm WT-2 | 551 | 141.75 | 386.01 | 5.45 |
| Sb+Zm WT-3 | 551 | 134.55 | 365.66 | 5.44 |
| Sb+Zm WT-4 | 551 | 139.59 | 379.2 | 5.43 |
| Sb+Zm WT-5 | 551 | 145.01 | 395.13 | 5.45 |
| Sb+Zm WT-6 | 551 | 133.13 | 362.05 | 5.44 |
| Sb+Zm WT-7 | 551 | 132.19 | 363.32 | 5.50 |
| Sb+Zm WT-8 | 551 | 113.15 | 308.31 | 5.45 |
| Sb+Zm WT-9 | 551 | 118.94 | 322.57 | 5.42 |
| Sb+Zm WT-10 | 551 | 127.65 | 349.83 | 5.48 |
| Sb+Zm WT-11 | 551 | 137.39 | 373.96 | 5.44 |
| Sb+Zm WT-12 | 551 | 141.31 | 387.02 | 5.48 |
| Sb+Zm WT-13 | 551 | 131.18 | 353.87 | 5.40 |
| Mean value |  |  | *Norm. factor*<sub>(sorghum)</sub> | **5.45** |

**Supplemental Table 6.** Ploidy calculation for *VuTAM2* knockout T2 lines.

| Sample ID | Gain | Test_peak (PK1<br>cowpea mean-x) | Ref_peak (PK2<br>soybean mean-x) | Test_peak/<br>Ref_peak | Ploidy |
| --- | --- | --- | --- | --- | --- |
| G23-TAM2_4-1_S1 | 551 | 191.44 | 319.67 | 0.60 | 2.00 |
| G23-TAM2_4-1_S2 | 551 | 184.37 | 306.47 | 0.60 | 2.01 |
| G23-TAM2_4-1_S3 | 551 | 187.63 | 317.73 | 0.59 | 1.97 |
| G23-TAM2_4-1_S4 | 551 | 186.28 | 310.58 | 0.60 | 2.00 |
| G23-TAM2_4-1_S5 | 551 | 197.41 | 320.68 | 0.62 | 2.05 |
| G23-TAM2_4-1_S6 | 551 | 194.52 | 321.00 | 0.61 | 2.02 |
| G23-TAM2_4-1_S7 | 551 | 196.07 | 321.77 | 0.61 | 2.03 |
| G23-TAM2_4-1_S8 | 551 | 191.94 | 320.28 | 0.60 | 2.00 |
| G23-TAM2_4-1_S9 | 551 | 189.19 | 312.15 | 0.61 | 2.02 |
| G23-TAM2_4-1_S10 | 551 | 197.89 | 325.30 | 0.61 | 2.03 |
| G23-TAM2_30-14_S1 | 551 | 187.24 | 311.81 | 0.60 | 2.00 |
| G23-TAM2_30-14_S2 | 551 | 190.66 | 314.53 | 0.61 | 2.02 |
| G23-TAM2_30-14_S4 | 551 | 186.38 | 315.54 | 0.59 | 1.97 |
| G23-TAM2_30-14_S5 | 551 | 184.10 | 308.65 | 0.60 | 1.99 |
| G23-TAM2_30-14_S6 | 551 | 179.38 | 304.06 | 0.59 | 1.97 |
| G23-TAM2_30-14_S9 | 551 | 182.95 | 313.87 | 0.58 | 1.95 |
| G23-TAM2_30-14_S10 | 551 | 182.34 | 302.98 | 0.60 | 2.01 |
| G23-TAM2_30-15_S1 | 551 | 182.23 | 304.76 | 0.60 | 2.00 |
| G23-TAM2_30-15_S2 | 551 | 186.90 | 312.10 | 0.60 | 2.00 |
| G23-TAM2_30-15_S3 | 551 | 192.21 | 321.89 | 0.60 | 1.99 |
| G23-TAM2_30-15_S5 | 551 | 181.31 | 301.20 | 0.60 | 2.01 |
| G23-TAM2_30-15_S6 | 551 | 188.03 | 312.11 | 0.60 | 2.01 |
| G23-TAM2_30-15_S8 | 551 | 171.52 | 289.10 | 0.59 | 1.98 |
| G23-TAM2_30-15_S9 | 551 | 180.39 | 300.86 | 0.60 | 2.00 |

